# Sensory neurons shape γδ T cell effector programs to control Psoriasiform Inflammation

**DOI:** 10.64898/2026.07.03.736363

**Authors:** Juan M. Inclan-Rico, Camila M. Napuri, Adriana Stephenson, Heather L. Rossi, Ulrich M. Femoe, Fungai Musaigwa, Li-Yin Hung, Huasheng Yu, Wenqin Luo, De’Broski R. Herbert

**Affiliations:** Department of Pathobiology, School of Veterinary Medicine, University of Pennsylvania, Philadelphia, Pennsylvania, USA; Department of Neuroscience, School of Medicine, University of Pennsylvania, Philadelphia, Pennsylvania, USA; Department of Microbiology and Immunology, School of Medicine, Tulane University, New Orleans, Louisiana, USA

**Keywords:** Neuroimmunology, Psoriasis, Skin, gamma delta T cells

## Abstract

Psoriasis is a chronic autoimmune skin disorder marked by IL-17 producing gamma delta T cell (γδT17) and pruritus, but immunoregulatory roles of itch-inducing neurons in this context remain unclear. This study addressed whether non-peptidergic (NP) afferents bearing the Mas-related G protein-coupled receptor D (MrgprD/NP1) and MrgprA3/NP2 subsets had differential effects on psoriasiform immunopathology. Data show human NP1 and NP2 neurons basally expressed an array of pattern recognition and cytokine receptor genes and psoriatic human skin had a profound dysregulation of neuropeptides and their receptors. In mice, imiquimod (IMQ) application reduced the density of MrgprD+ skin afferents, whereas NP1 neuron ablation exacerbated IMQ-induced disease. Strikingly, NP1 activation using either optogenetics or β-alanine before IMQ exposure significantly reduced epidermal thickness, psoriatic clinical score and γδT17 cell accumulation. In stark contrast, NP2 activation increased the numbers of γδT17 cells that co-expressed amphiregulin (Areg) and exacerbated IMQ-driven skin pathology. Instead, pre-emptive NP1 stimulation shifted γδ T cell profiles away from being IL-17 and Areg dominant to IL-13+ γδ T cells expressing the transcription factor GATA3 accompanied by IL-10 secretion. Importantly, IL-10 signaling blockade reversed NP1-mediated suppression of IMQ-induced dermatitis. These data show that inflammatory skin disease can be distinctly modulated by sensory neuron subsets.

## Introduction

Psoriasis is a chronic and debilitating autoimmune disorder that affects 2-3% of the global population, including >7.5 million adults in the U.S, with the highest incidence occurring between the ages of 50-59 (Armstrong et al., 2021, Armstrong and Read, 2020). Concerningly, >40% of psoriatic patients report a significant negative effect on their quality of life and their ability to work (Wang et al., 2024). Despite being a heterogeneous disease, plaque psoriasis, characterized by the development in the skin of well-demarcated dry and red plaques with silver scale, is the most prevalent clinical manifestation, accounting for 70-80% of cases (Armstrong et al., 2021, Armstrong and Read, 2020). Psoriatic plaque formation is triggered by aberrant responses against endogenous molecules such as nucleic acids and antimicrobial peptides recognized by pattern recognition receptors (PRR) expressed in plasmacytoid dendritic cells and keratinocytes (Armstrong et al., 2025). PRR signaling in these cells induces their rapid secretion of type 1 interferons and TNF, which in turn stimulate antigen-presenting cells (APC) to release cytokines like IL-1β, IL-6, and IL-23. These signals activate innate γδ T cells and promote the differentiation of T_H_17 cells that together robustly secrete IL-17 cytokines (IL-17A/F, and IL-22) that promote keratinocyte hyperplasia and proliferation (Armstrong et al., 2025). While immunotherapies that inhibit TNF, IL-23, IL-17, and other cytokines have proven very successful in improving the quality of life of patients living with psoriasis (Guo Jia et al., 2023), these therapies can lead to disease aggravation or increased risk of infections, highlighting the need for a clearer understanding of how psoriatic inflammation is regulated.

Clinically significant itch or pruritus is highly prevalent in 64-98% psoriatic patients (Junsuwan et al., 2023). Itch is mainly transmitted by sensory nerve afferents that innervate the epidermis and dermis and transmit these signals to their cell bodies emanating from dorsal root ganglia (DRG) or trigeminal ganglia (TG). Recent single-soma deep sequencing analysis of human DRG neurons revealed three molecularly distinct subsets of non-peptidergic (NP) sensory neurons, namely NP1, NP2, and NP3; that likely function to transmit itch and are remarkably conserved between humans and mice (Usoskin et al., 2015, Yu et al., 2024). NP1 and NP2 neurons distinctively express specific Mas-related G-protein coupled receptors (MRGPR), including *MRGPRX1* (shared by NP1 and NP2), *MRGPRD* (enriched only in NP1), *MRGPRX3,* and *MRGPRX4* (unique to NP2) (Han et al., 2013, Liu et al., 2012, Liu et al., 2009). Conversely, NP3 neurons have low MRGPR levels but distinctively express the neuropeptide SST. Existing evidence suggests that NP1-3 afferents are polymodal, with sensitivity to mechanical (NP1 and NP2), chemical (NP1-NP3), and thermal stimuli (NP1-NP3) (Beattie et al., 2022, Cranfill, 2025, Guo Changxiong et al., 2023, Zylka et al., 2005). We and others have demonstrated that these itch-transmitting afferents can modulate cutaneous immunity (Deng et al., 2023, Fassett et al., Inclan-Rico et al., 2024, Inclan-Rico et al., 2025, Liu et al., 2025, Riol-Blanco et al., 2014, Zhang et al., 2021). While mouse NP2 neurons appear to mostly trigger pro-inflammatory responses(Inclan-Rico et al., 2024, Liu et al., 2025), NP1 (MrgprD+) neurons were recently shown to suppress skin anti-bacterial immunity(Zhang et al., 2021). Nevertheless, how discrete itch-inducing afferents contribute to the immunopathology of psoriasis is undetermined.

Gamma delta (γδ) T cells are a major lymphocyte population in skin and are increasingly recognized as key regulators of barrier immunity (Hu et al., 2023). In mice, γδ T cell populations are often categorized according to their T cell receptor (TCR) repertoire and cytokine-producing potential, with Vγ4⁺ and Vγ6⁺ subsets serving as major sources of IL-17 during psoriasiform inflammation (Cai et al., 2012, Ramirez-Valle et al., 2015). Despite intense focus on the pathogenic role(s) for IL-17+ γδ T cells (γδT17) subsets, emerging evidence indicates γδ T cells are functionally heterogeneous and secrete immunoregulatory cytokines such as IL-10 and IL-13 (Peters et al., 2018). However, it remains entirely unknown whether tissue-derived signals from cutaneous sensory neurons can influence pro-inflammatory vs. regulatory γδ T cell states during inflammatory skin diseases.

This study shows that human NP1 and NP2 neurons selectively expressed high levels of PRRs and cytokine receptors linked to IL-17-associated immunopathology. Moreover, transcriptional analysis of human psoriatic lesional and non-lesional skin revealed dysregulated expression of multiple neuropeptides, and neuropeptide receptors during disease. Analysis of murine psoriasiform skin revealed that γδT17 cells express many neuropeptide receptors notably, glutamate receptor expression, supporting the possibility that NP1 neuron-derived mediators may regulate pathogenic γδ T cell function. Using the TLR7 agonist Imiquimod to induce murine psoriasiform immunopathology (IMQ)(Que et al., 2025), data show that basal epidermal NP1 innervation was reduced and genetic depletion of NP1 neurons or activation of NP2 neurons prior to IMQ application exacerbated IL-17-associated skin immunopathology. On the other hand, activation of NP1 neurons using optogenetics or the MrgprD ligand, β-alanine, blunted psoriasiform immunopathology. Congruently, NP1 neuron activation caused a reduction in secretion of IL-17 associated cytokines and reduced skin neutrophils. Multiparametric flow cytometric analysis revealed that IMQ treatment augmented γδ T cells expressing IL-17, Areg, and Ki-67, but that NP1 neuron stimulation prior to IMQ treatment significantly reduced these effects. Instead, pre-emptive NP1 activation increased IL-13+ γδ T cells with GATA3 expression and elevated the release of IL-13 and IL-10 by sdLNs, suggesting a shift in γδ T subset activation via NP1 afferents. Together, these results support a model wherein loss of NP1 afferents during IMQ-induced dermatitis contributes to IL-17-mediated inflammation by shifting γδ T cell phenotypes.

## Materials and methods

### Study design

Mouse experiment group sizes ranged between four and six mice (matched for age and sex) and were repeated two to three times to ensure reproducibility. Mice were tattooed for identification, with experimental groups randomized across cages to account for any microisolator effects. To address subjectivity during the study, animal cages and experimental samples/groups were assigned a letter number code to ensure that experiments were conducted in a blinded manner. For all flow cytometry experiments, negative controls were included, such as fluorescence minus one controls, to establish reliable and reproducible gates for each marker.

### Mice and experimental procedures

*Mrgprd^tm1.1(cre/ERT2)Wql^*/J (MrgprD^CreERT2^) (Olson et al., 2017) were generated and maintained in-house. Tg(Mrgpra3-GFP/cre)#Xzd (MrgprA3 Cre) mice were originally provided by Xinzhong Dong (Johns Hopkins University). B6.Cg-*Gt(ROSA)26Sor^tm9(CAG-tdTomato)Hze^*/J (Ai9 mice) (Madisen et al., 2010), B6.Cg-Gt(ROSA)^26Sortm32(CAG-COP4*H134R/EYFP)Hze^/J (Rosa^ChR2-EYFPf/f^) (Nagel et al., 2003), B6.129P2-Gt(ROSA)26Sor^tm1(DTA)Lky/J^ (ROSA-DTA) and wildtype mice (C57BL/6J) strains were purchased from Jackson Laboratories. MrgprD- or MrgprA3-tdTomato mice were generated by interbreeding Mrgprd^CreERT2^ or MrgprA3^Cre^ and Ai9 mice, whereas MrgprD-DTA mice were raised by crossing Mrgprd^CreERT2^ and ROSA-DTA mice. MrgprD-ChR2 or MrgprA3-ChR2 mice were generated by crossing Mrgprd^CreERT2^ or MrgprA3^Cre^ and Ai32 mice. Optogenetic and ablation strains were bred to heterozygosity and compared with age- and gender-matched controls from a separate set of breeders housed in the same facility. Co-housed Cre-negative littermate controls were used for all optogenetic and neuronal ablation experiments. Only 8–10-week-old male mouse cohorts were used in all experiments of this manuscript. All mice were housed under specific pathogen-free conditions in 12/12h light/dark cycles with controlled temperature at ±22.2°C and humidity ±20% of 50%. at the University of Pennsylvania. 4-6-week-old MrgprD-tdTomato, MrgprD-DTA, or MrgprD-ChR2 mice and their respective controls were treated via i.p. with 0.5 mg of tamoxifen (extracted in sunflower seed oil) for seven consecutive days, followed by an additional 14 days of rest to drive gene expression. All procedures were approved by the Institutional Animal Care and Use Committee of the University of Pennsylvania (protocol 805911).

For induction of psoriasiform inflammation, mice were treated with 62.5 mg of Vaseline (as negative control) or 5% imiquimod (IMQ) cream from 0.25 g single-use packets on ventral and dorsal sides of the ear pinnae for five or seven consecutive days. Ear thickness was assessed using calipers, taking the average of 3 consecutive measurements as the value for each mouse. Treatment with IMQ was completed after all other assessments (e.g. optogenetic stimulations or caliper measurements) were completed. For β-alanine intradermal injections, mice were anesthetized with 4% isoflurane and injected intradermally into the ear pinnae with 20 μL PBS or 50mM beta-alanine, every 2 days for 5 days. For cytokine neutralization experiments, mice were treated intraperitoneally (i.p.) with 750 µg of neutralizing antibody against IL-10R (Bioxcell, cat# BE0050), three and one day before IMQ treatment. Mice were euthanized by CO_2_ for all tissue recovery procedures following American Veterinary Medicine Association guidelines

### Optogenetic stimulation

Mice were anesthetized with 1-4% isoflurane to prevent facial movement and scratching and maintained on a heating pad during photostimulation. Optogenetic stimulation was performed using a 473nm 100mW diode-pumped solid state (DPSS) laser (SLOC Lasers, Shanghai, China), with an attached power supply. An optical cannula aimed at the dorsal ear was placed 1-2 cm from the surface of the skin. Stimulation was performed with a sinus waveform generator to pulse 10Hz, pulse length for 30min with a power density of 8-10 mW/mm^2^. Mice underwent five 30-minute stimulation daily sessions.

### Skin and skin-draining lymph node preparation for flow cytometric analysis

Ear skin explants were surgically dissected and processed using previously established laboratory protocols. Skin biopsies were mechanically dissociated and then incubated with skin digest solution containing 2mg/ml of collagenase XI (Sigma-Aldrich), 0.5mg/ml hyaluronidase (Sigma-Aldrich), and 0.1 mg/ml DNase (Roche) dissolved in DMEM containing 2.5% FBS for 25min at 37°C with constant agitation. Skin homogenates were then passed through an 18-gauge needle 3-5 times and incubated at 37°C for an additional 20min. Then, cell preparations were passed again through an 18-gauge needle and filtered twice through 100mm and 40mm filters. Fresh cells were suspended in 10% FBS for enumeration and stained for flow cytometry as indicated below. Skin-draining lymph nodes were mechanically dissociated in 1% FBS DMEM, passed through 70uM filters, spun down for 5 minutes at 1700 RPM, and supernatant removed. Cells were then resuspended in RPMI supplemented with 10% FBS and incubated with Cell Activation Cocktail (with Brefeldin A) for 5-6h for measurement of intracellular cytokine expression.

Skin and sdLN cell suspensions were stained for live/dead cell exclusion using LIVE/DEAD™ Fixable Aqua Dead Cell Stain Kit according to manufacturer’s protocol followed by Fc Block for 15min at 4°C, and surface marker staining for 25 min at 4°C. Then, eBioscience™ Foxp3/Transcription Factor Staining Buffer Set was used according to manufacturer’s protocol. Antibodies for nuclear proteins were incubated overnight at 4°C. Intracellular staining was done the next day for 1-1.5hrs on ice. Cells were analyzed with a BD Symphony A3 Lite using the BD FACSDiva 9.0 Software. All antibodies are listed in Supplementary Table 1.

### Immunofluorescence

Skin tissue samples collected were fixed in cold 4% PFA for 2 hrs at 22°C, washed with 1XPBS for 1- 4 hrs, and left in 30% sucrose (in 1x PBS) overnight 4°C. Samples were then embedded with OCT, and 20-µm-thick sections were cryosectioned. Sections were re-fixed in cold 4% PFA for 15 min. and incubated for 90 min at 22°C in blocking buffer (1X TBS, 1% BSA, 5% normal donkey serum, Fc block, 0.3% Triton X-100). After rinsing three times with 1XTBS 1% BSA, sections were incubated in primary antibodies diluted in blocking buffer at 4°C overnight. Tissue sections were rinsed and incubated with secondary antibodies diluted in blocking buffer for 120 min 22°C., incubated with DAPI diluted in 1X TBS 1%BSA for 10 min. at 22°C, mounted on glass slides with fluoroshield mounting medium before imaging. The primary antibodies used for IF staining were chicken anti-mouse Keratin 5 Polyclonal Antibody (1:500 dilution; BioLegend), rabbit anti-cytokeratin-10 antibody (EP1607IHCY; 1:500 dilution; Abcam), biotinylated mouse anti-mouse beta tubulin 3 (TUBB3) antibody (1:500 dilution; BioLegend), Armenian hamster anti-mouse TCR γ/δ Antibody (1:500 dilution; BioLegend), rat anti-mouse MHC Class II antibody [M5/114] (1/100; Abcam); and DAPI solution (1:1000 dilution; Thermo Fisher). The secondary antibodies used were AF488 streptavidin, AF488 Armenian hamster, AF647 Donkey anti-rat, AF488 Donkey anti-chicken, all at 1:500 dilution. Exposure times and fluorescence intensities were normalized to appropriate control images per channel separately and then merged. Epidermal thickening was measured using the ‘scale bar’ function of the Leica Application Suite X version 5.1.0.25446, drawing a straight line from the top stratum corneum layer to the base stratum basale layer. Five measurements per biological replicate were taken for an average total measurement. Epidermal nerve density was measured using Image J similar to previously described (Valero-Pacheco et al., 2022). After splitting each image into individual channels and adjusting the threshold using the Otsu method, eight equally-sized regions of interest (ROIs) were defined in the mid-section of the epidermis using the ‘Square’ selection tool. Then, the percent (%) area for each ROI was calculated, and the eight ROI measurements per image were averaged. At least three representative pictures per biological replicate were evaluated independently.

### Histology and Pathology Scoring

Skin sections were fixed in 10% formalin buffered for at least 24 hrs., followed by paraffin-embedding using a Leica modular tissue embedding system and sliced into 7-mm sections using a Leica RM2235 rotatory microtome (Leica Biosystems). Tissue sections were deparaffinized, hydrated, and stained with hematoxylin and eosin (H&E). Images were acquired with a Leica DM6000 Upright Widefield Fluorescence Microscope and Nikon Eclipse Ti2-E inverted microscope, with epidermal thickness measured as described above. Clinical scoring was performed by a blinded dermatopathologist and assigned a numeric value using the following scale 0 – No significant findings, 1 – Minimal, 2 – Mild, 3 – Moderate, 4 – Severe. Heatmaps were generated of averaged scores from biological replicates of the same group.

### ELISA

Cervical lymph node tissue was dissected and processed using DMEM-F12 containing 5% FBS. Tissue homogenates were filtered through a 70uM cell strainer and cell suspensions spun down for 5 min. at 1700 RPM in 4°C centrifuge. Samples were plated on a 96-well u-bottom plate with CD3/CD28 activation cocktail for 72 hrs. and stored in -20°C. Commercially available assays listed in Supplementary Table 1 were used to measure cytokine concentration levels.

### *In silico* analysis of publicly-available single-cell RNA sequencing

Datasets from GSE173706 and GSA: HRA011011 (Chen et al., 2025, Ma et al., 2023) were further processed with Seurat 4.0 R package(Hao et al., 2021). For quality control, only genes expressed in at least 3 cells and cells expressing at least 200 genes were included. Cells expressing >10% mitochondrial genes were excluded from the downstream analysis. Data were normalized and scale using default parameters and the number of principal components were estimated using *RunPCA* followed by *ElbowPlot*. Uniform Manifold Approximation and Projection (UMAP) was used for dimensionality reduction and performed using *RunUMAP*. Markers for cell clusters were identified using the *FindAllMarkers* function equipped in the Seurat and cell types were annotated manually using canonical markers. To further resolve cell subtypes (e.g., B cells), the subset function in Seurat was used, with subset of cells subjected to a second round of principal components identification and dimensional reduction as the same as above. We deployed the processed results in Shinny App for further exploration.

### Statistics

Results are shown as the mean ± s.e.m. P < 0.05 was considered as significantly different. Grubb’s test was used to identify outliers. Normal distribution of the data was determined by Shapiro-Wilk test. When these data were normally distributed, we used parametric statistical tests; otherwise, non-parametric analysis was used when stated. Statistical analysis was performed using Student’s t-test for two groups, one-way ANOVA for three groups, or two-way ANOVA (ear thickness measurements), with appropriate post-hoc tests (Bonferroni for One-Way and Tukey’s for Two-Way ANOVA in Prism 11.0.2 (100) (GraphPad Software).

## Results

### Human NP neurons are enriched for receptors associated with psoriasis

Psoriasis is characterized by the formation of scaly plaques driven by excessive keratinocyte proliferation and maturation initiated by the aberrant activation of pattern recognition receptors (PRR) leading to the release of IL-17 cytokines (de Koning et al., 2010, Lande et al., 2007, Shallev et al., 2018). Psoriatic patients commonly experience intense itch or pruritus (Junsuwan et al., 2023); however, the contributions of specific subsets of itch-inducing sensory neurons to the pathology of psoriasis are poorly understood. In mice and humans, three main subsets of non-peptidergic neurons, namely NP1, NP2, and NP3, have been identified to transmit chemical-induced itch (Usoskin et al., 2015, Yu et al., 2024). We hypothesized that itch-evoking neurons could be engaged during psoriasis, potentially through receptor signaling for soluble mediators released during disease. To evaluate this idea, we analyzed PRR and cytokine receptor expression levels in NP neuron subsets within human DRG using our previously published deep RNA-sequencing data (Yu et al., 2024). Consistent with our hypothesis, human NP1 neurons were highly enriched for PRR-coding genes strongly associated with psoriasis susceptibility, including nucleic acid sensors (*TLR7*, *DDX58* (RIG-I), and *CGAS*), inflammasome components (*NLRP1,3,10*; *NOD2, PYCARD, CARD6*), and C-type lectins (*CLEC2B, 7A*) (**Fig. 1A,B**). NP2 neurons expressed TLR4, TLR8, and IFIH1, the latter of which encodes the cytosolic receptor MDA5, all of which are associated with susceptibility or disease progression of psoriasis (Coto-Segura et al., 2023, Kim et al., 2016, Smith et al., 2016). Moreover, NP1 neurons from healthy human donors showed high expression of the receptors for IL-17, IL-1β, IL-23, TNF, and IFNI/II, whereas NP2 nerves were enriched for IL-1β, IL-6, and TNF receptors (**Fig. 1C**).

**Figure 1.**
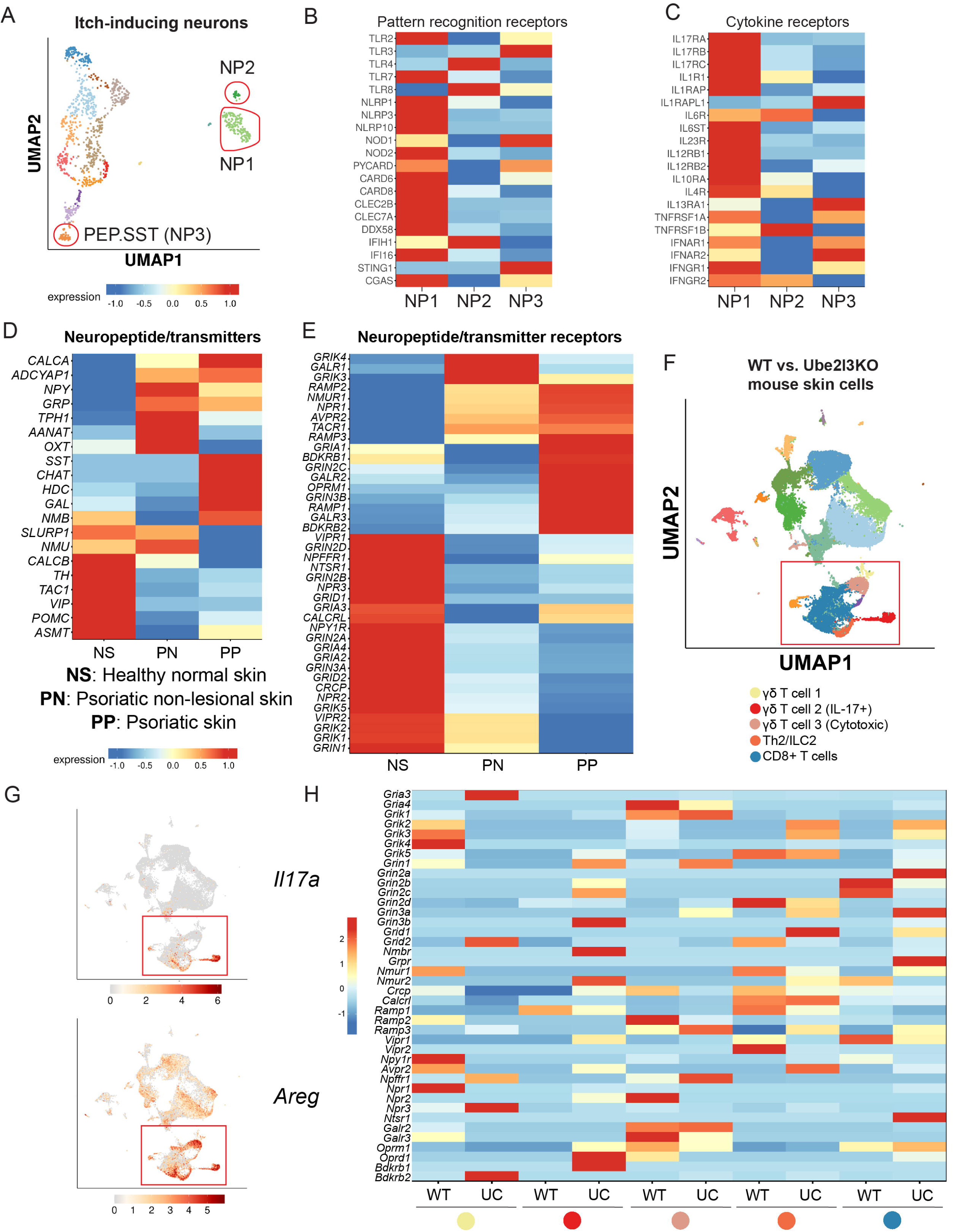
Transcriptional analysis of neuro-immune pathways related to psoriasis. (**A**), Uniform Manifold Approximation and Projection (UMAP) plot illustrating defined cell clusters generated from deep-sequencing of human DRG neurons (Yu, *et al*. 2024). Squares highlight clusters of non-peptidergic (NP) neurons. (**B**), Heatmap showing expression of pattern recognition receptors (PRR) or (**C**), cytokine receptors in NP neuron clusters from A. (**D**), Heatmaps showing publicly-available (GSE173706, Ma, *et al*. 2023) transcriptional data of neuropeptides, neurotransmitter-synthesizing enzymes, and (**E**) their receptors in healthy human skin (NS), non-lesional biopsies (PN), and active psoriatic lesions (PP). (**F**), UMAP plot illustrating cell clusters defined by single-cell RNA sequencing of murine skin populations from keratinocyte-specific Ube2l3-deficient mice that resemble human psoriasis (GSA: HRA011011, Chen, *et al*. 2025). Square highlights lymphocyte clusters. (**G**), *Il17a* and *Areg* expression from F. (**H**), Heatmaps of neuropeptide and neurotransmitter receptors in lymphocyte clusters from F.

### Altered expression of neuronal soluble mediators and their receptors in psoriatic skin

We then evaluated publicly-available data from healthy skin, non-lesional and lesional psoriasis skin for RNA expression levels of neuron-derived soluble mediators, including neuropeptides, neurotransmitters, and their respective receptors (Ma et al., 2023). This *in silico* transcriptomic analysis showed an enrichment of neuropeptides in psoriatic lesional and non-lesional skin, including CGRP, PACAP, NPY, and GRP compared to healthy skin. SST, GAL, and NMB were also elevated in psoriatic skin (**Fig. 1D**). Moreover, while receptor expression for NMU, CGRP, and Substance P was enriched in psoriatic lesions, receptors for VIP, Neuropeptide FF, and glutamate were curiously downregulated (**Fig. 1E**), suggesting a marked dysregulation of neuron-derived mediators in patients with active psoriasis. Similarly, in a mouse model where genetic conditional ablation of Ube2l3 in keratinocytes results in psoriasis-like pathology (Chen et al., 2025), there was increased expression of receptors for CGRP, Nmb, VIP, endogenous opioids, bradykinin, and glutamate on γδ T cell populations that show enriched IL-17 and Areg expression (**Fig. 1F-H**). Other cutaneous lymphocyte subsets, such as Th2, ILC2s, and CD8+ T cells, also presented with differences in receptor expression. Altogether, these analyses suggested that PRR and cytokine receptors expressed in skin-innervating afferents of NP neurons could potentially allow crosstalk with immune cells in the same microenvironment through soluble mediators they may release during psoriasis.

### Loss of MrgprD+ afferent innervation following IMQ treatment

To investigate potential communication between neurons and immune cells, we employed an established murine model for inducing psoriasiform inflammation. Repetitive application of 5% IMQ cream on the ear pinnae of C57BL/6 mice induces psoriasiform features of human pathology, such as skin thickening, scaling, acanthosis, and hyperkeratosis (van der Fits et al., 2009). As expected, 5 days of IMQ application resulted in elevated Krt5 expression in basal and suprabasal epidermal cells and Krt10 staining in suprabasal keratinocytes as well as increased epidermal thickness compared to Vaseline-treated controls (**Fig. 2A-F**). Next, we tested whether IMQ exposure had any impact on skin neuronal innervation. Immunofluorescence staining for the pan-neuronal marker βIII-tubulin revealed that the percentage area covered by βIII-tubulin+ innervation in the epidermis was significantly reduced by IMQ-treatment (**Fig. 2G-I**). NP1 neurons distinctly express MrgprD and it was reported that IMQ-exposure in mice reduced MrgprD mRNA expression levels in DRG neurons (Sakai et al., 2016). To specifically identify NP1 skin afferents, MrgprD-tdTomato reporter mice were generated by interbreeding *Mrgprd^CreERT2^* (Olson et al., 2017) and *Ai9* (Madisen et al., 2010) mouse strains. Co-localization of MrgprD-tdTomato+ fibers with βIII-tubulin+ nerve fibers confirmed visualization of skin NP1 nerve afferents in the ear pinnae (**Fig. S1A**). MrgprD-tdTomato+ afferents densely innervated the epidermis of Vaseline-treated mice as previously shown (Zylka et al., 2005) (**Fig. 2J**) but, unexpectedly, we found a substantial reduction in this pattern following IMQ treatment (**Fig. 2K,L**). Moreover, skin-resident myeloid and lymphoid subsets are known to co-localize with cutaneous nerve afferents (Enamorado et al., 2023, Hanč et al., Inclan-Rico et al., 2024). Thus, we quantified the distance of γδ T cells or MHC-II+ cells to MrgprD+ skin afferents in Vaseline- vs. IMQ-treated MrgprD-tdTomato reporter mice via immunostaining. In Vaseline-treated mice, epidermal γδ T cells were located proximal to tdTomato+ NP1 afferents; however, IMQ exposure increased the distance between these cells (**Fig. 2M-O**). MHC-II+ cells were mostly located in the dermis of Vaseline-exposed mice and IMQ treatment further separated them from epidermal MrgprD-tdTomato+ nerves (**Fig. S1B-D**). Overall, these data indicate that MrgprD+ afferents densely innervate the epidermis and co-localize close to γδ T cells under basal conditions, but IMQ exposure reduces epidermal MrgprD+ innervation and increases their distance from γδ T cells.

**Figure 2.**
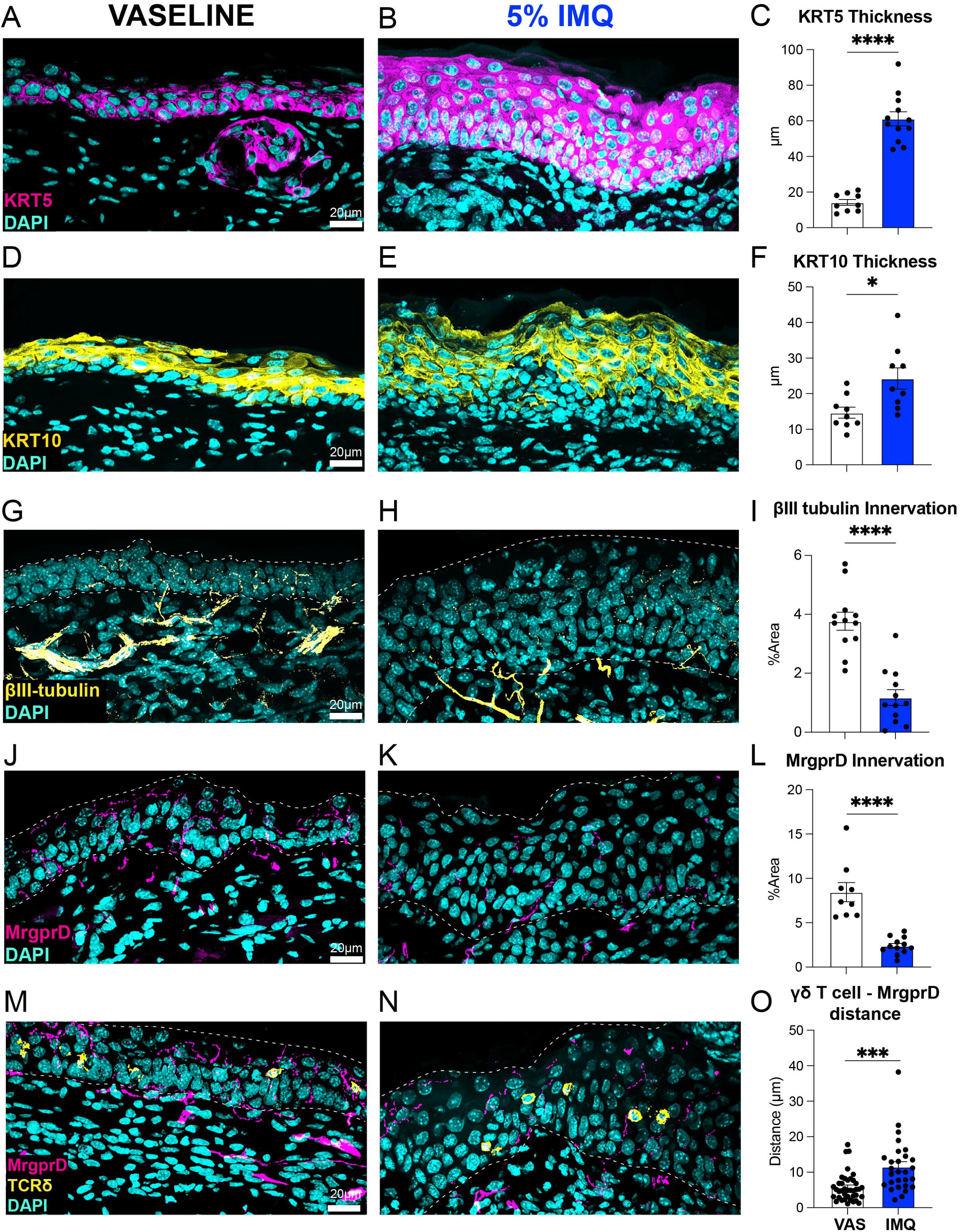
**Keratinocyte and neuronal innervation changes induced by IMQ application.** (**A,B**), Immunofluorescence (IFA) staining for Keratin-5 (Krt5), or (**D,E**), Keratin-10 (Krt10) of ear skin sections from MrgprD-tdTomato reporter mice treated with Vaseline or 5% IMQ cream for 5 consecutive days. (**C,F**), Quantification of epidermal thickness measured from the basal layer until the end of Krt5 or Krt10 staining in the epidermis. (**G,H**), IFA staining with the pan-neuronal marker βIII-tubulin, (**J,K**), epidermal MrgprD-tdTomato+ afferents, or (**M,N**), TCRδ+ cells (yellow) and MrgprD-tdTomato+ nerve afferents in Vaseline- or IMQ-treated mice. (**I,L,O**), Mean ± s.e.m (n=9-12) of the percentage Area covered by βIII-tubulin+ or MrgprD-tdTomato+ nerve afferents or (n=28-39) the distance between γδ T cells and MrgprD+ nerve afferents in the epidermis. *P* values were determined by two-tailed Student’s t-tests *P<0.05, **P<0.01, ***P<0.001, ****P<0.0001. Representative of 2 independent experiments, each with ≥3 biological replicates.

### Ablation of MrgprD+ neurons exacerbates IMQ-induced dermatitis

Several studies recently defined how itch-transmitting neurons can either promote or suppress skin inflammation (Deng et al., 2023, Fassett et al., Inclan-Rico et al., 2024, Inclan-Rico et al., 2025, Liu et al., 2025, Zhang et al., 2021). While NP2 and other TRPV1+ neurons promote IL-17-mediated responses during psoriasis or against parasites and fungi, NP1 neurons can inhibit anti-bacterial responses (Deng et al., 2023, Fassett et al., Inclan-Rico et al., 2024, Inclan-Rico et al., 2025, Liu et al., 2025, Riol-Blanco et al., 2014, Zhang et al., 2021). Based on the reduced epidermal density of MrgprD+ afferents upon IMQ exposure, we predicted that their ablation may exacerbate IMQ-induced psoriasiform pathology. As a loss-of-function approach, we generated *MrgprD^CreERT2^:Rosa^DTA^* (MrgprD-DTA) transgenic mice. In this system (Cranfill et al., 2025), tamoxifen administration induces Cre-mediated recombination in MrgprD-expressing neurons, removing a loxP-STOP-loxP cassette that otherwise blocks diphtheria toxin A (DTA) expression allowing NP1 neuronal ablation. Two weeks after tamoxifen injection, control and MrgprD-DTA mice exposed to IMQ had substantial epidermal thickening that was significantly higher in mice depleted of MrgprD+ neurons, as shown by H&E staining (**Fig. 3A,B**). Clinical scoring showed increased signs of IMQ-induced psoriasiform dermatitis in MrgprD-DTA mice, including hyperkeratosis, subcorneal pustules, basal hyperplasia, and neutrophil and lymphocyte infiltration (**Fig. 3C**). Consistently, flow cytometric analysis showed elevated percentages and numbers of skin neutrophils (CD45^+^ CD11b^+^ Ly6G^+^ Ly6C^mid^) in MrgprD+ neuron depleted mice exposed to IMQ (**Fig. 3D,E**). Keratinocyte hyperplasia and neutrophil recruitment during IMQ-triggered dermatitis is largely driven by IL-17 cytokines (IL-17A/F) and IL-22 (Furue et al., 2020); thus, production of these cytokines was measured following culture of skin-draining lymph nodes (sdLNs). Restimulated (αCD3/αCD28) sdLNs from MrgprD-DTA mice produced significantly higher levels of IL-17A, IL-17F, and IL-22 than control mice (**Fig. 3F**). Since γδ T cells are recognized as key drivers of IL-17 inflammation during IMQ-induced disease (Cai et al., 2011), we characterized cytokine/cytokine receptor and proliferation profiles of γδ T cells using the self-organizing map (SOM) tool FlowSOM for unsupervised clustering and dimensionality reduction (Van Gassen et al., 2015). Data show that γδ T cells from IMQ-exposed mice had high levels of IL-17, Amphiregulin (Areg), and the proliferation marker (Ki-67) (**Fig. 3G-I**). While IL-17 and Areg were co-expressed in γδ T cells, those enriched for Ki-67 were moderately separated. Nonetheless, depletion of MrgprD+ neurons resulted in higher percentages and numbers of IL-17^+^Areg^+^ and Ki-67^+^ γδ T cells compared to controls (**Fig. 3J,K**). Further, unbiased multiparametric analysis revealed 5 distinct γδ T cell clusters, of which Clusters 1-3, enriched for IL-17, Areg, TNF, and Ki-67, were more abundant in MrgprD-DTA mice versus controls (**Fig. 3L-N**). These data revealed that depletion of MrgprD+ neurons increases IMQ-induced IL-17 responses and epidermal pathology.

**Figure 3.**
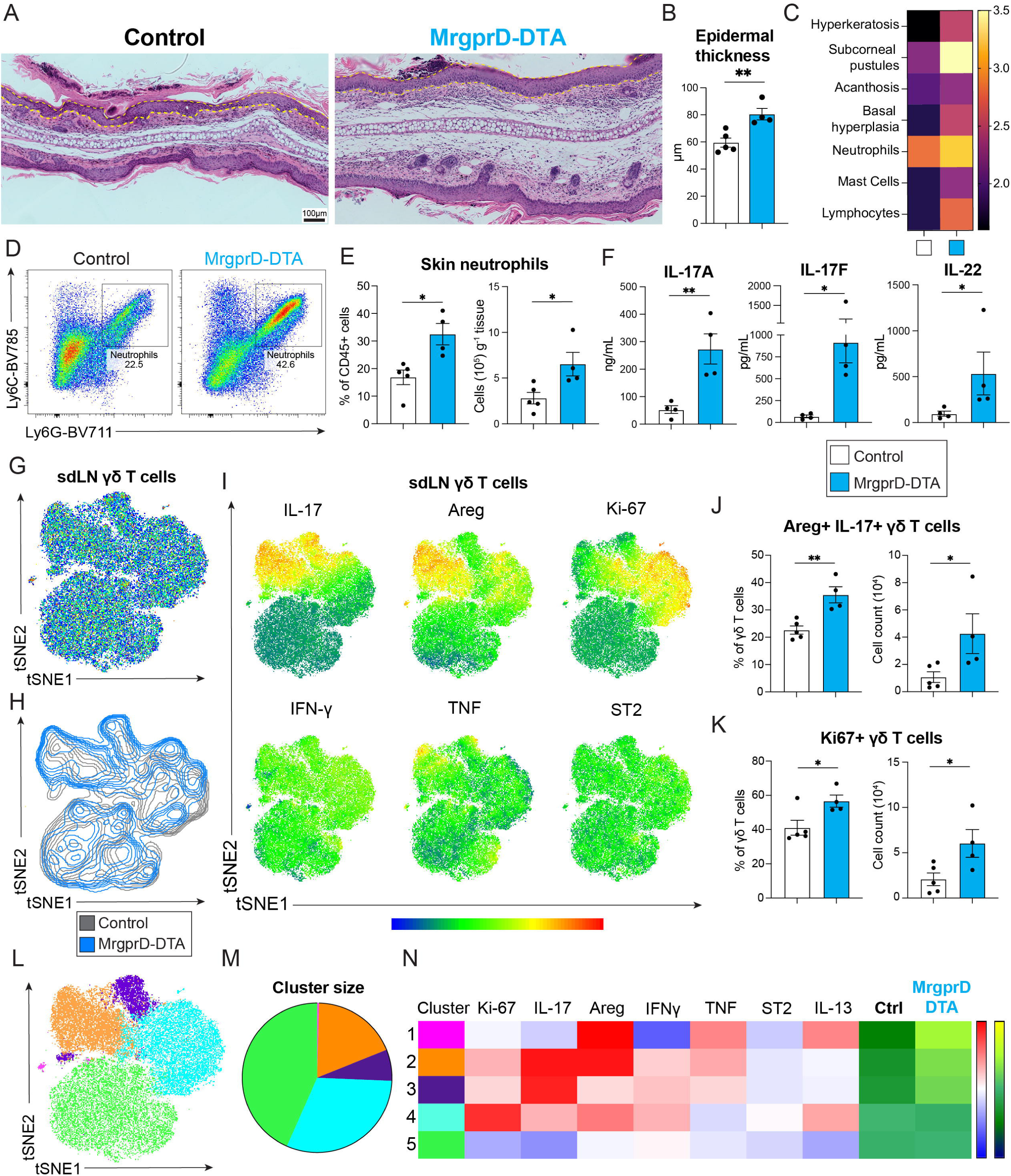
Ablation of NP1 neurons exacerbates IMQ-induced skin pathology and γδ T cell activation. (**A,B**), H&E staining and epidermal thickness quantification of FFPE ear skin sections from control or MrgprD-DTA mice after 7 days of treatment with 5% IMQ cream. Dotted lines indicate epidermal thickness. (**C**), Heatmap representing the averaged clinical score values of A. (**D,E**), Representative dot plots and quantification of skin neutrophils (CD45^+^ CD11b^+^ Ly6G^+^ Ly6C^int^) assessed by flow cytometry. (**F**), Concentration of IL-17A, IL-17F, and IL-22 in cell-free supernatants from skin-draining lymph nodes (sdLNs) stimulated with αCD3/αCD28 antibodies for 72hrs. (**G,H**), t-SNE dot plots and counterplots of concatenated γδ T cells from sdLNs of control or MrgprD-DTA mice after IMQ treatment. (**I**), Expression of cytokines, Ki67, and ST2 in t-SNE dot plots from G. (**J,K**), Percentage and absolute numbers of IL-17^+^ Areg^+^ or Ki-67^+^ γδ T cells quantified with traditional flow cytometric gating. (**L-N**), FlowSOM analysis, cluster sizes, and heatmaps of γδ T cell clusters from G. Bar graphs depict mean ± s.e.m (n=4-5). *P* values were determined by two-tailed Student’s t-tests. *P<0.05, **P<0.01. Representative of 2-3 independent experiments, each with ≥4 biological replicates.

### Stimulation of NP1 neurons reduces IMQ-induced skin inflammation

To further define NP1 neuron biology in the context of IMQ-induced dermatitis, we turned to a gain-of-function optogenetic approach to selectively activate MrgprD+ afferents *in vivo* using *MrgprD^CreERT2^:Rosa^ChR2-EYFPf/f^* (MrgprD-ChR2) transgenic mice that selectively express the light-sensitive non-selective cation channel channelrhodopsin 2 (ChR2) (Olson et al., 2017, Sharif et al., 2020). In this system, skin exposure to 473nm blue light opens ChR2, resulting in cation influx, action potentials, and the release of soluble mediators (Xin et al., 2023). Using established protocols (Cohen et al., 2019, Inclan-Rico et al., 2024), MrgprD^Cre^ negative ChR2 flox control (control) and MrgprD-ChR2 mice were exposed to blue light (10-12 mW, pulses of 30s and 10Hz) in the ear pinnae for 30 mins/day immediately before topical application of 5% IMQ cream or Vaseline (negative control) for 5 days (Que et al., 2025) (**Fig. 4A**). Strikingly, there was a significantly attenuation of IMQ-induced ear thickening in MrgprD-ChR2 mice between 3-5 days post-treatment compared to littermate controls (**Fig. 4B**). No increase in ear thickness occurred following Vaseline treatment of either strain.

**Figure 4.**
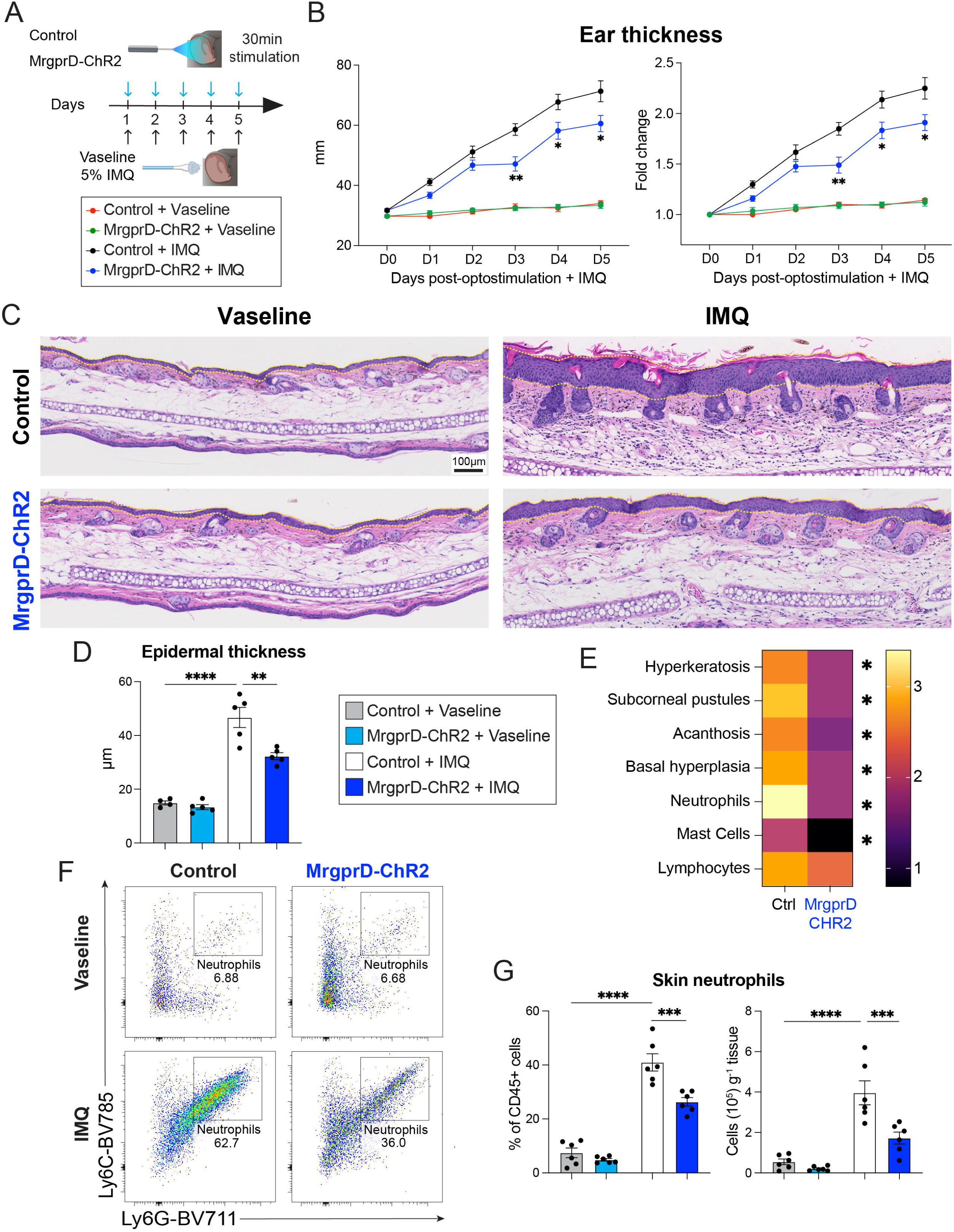
**NP1 neuron optogenetic activation limits the severity of IMQ-induced psoriatic pathology.** (**A**), Experimental approach for optogenetic ear stimulation of ChR2 control or MrgprD-ChR2 mice followed by topical application of Vaseline or 5% IMQ cream. (**B**), Raw values and fold change of ear thickness measured daily. (**C,D**), H&E staining and epidermal thickness quantification of FFPE ear skin sections from photostimulated control or MrgprD-ChR2 mice after 5 days of treatment with Vaseline or 5% IMQ cream. Dotted lines indicate epidermal thickness. (**E**) Heat map representing the average of clinical score values of IMQ-exposed ear sections analyzed as in C. (**F,G**), Representative dot plots, percentages, and absolute numbers of of skin neutrophils (CD45+ CD11b+ Ly6G+ Ly6Cint) assessed by flow cytometry. Line or bar graphs depict mean ± s.e.m (n=4-5). *P* values were determined by two-tailed Student’s t-tests, One-way ANOVA with Bonferroni post hoc correction, or Two-way ANOVA with Tukey’s for post hoc test. *P<0.05, **P<0.01, ***P<0.001, ****P<0.0001. Representative of 2-3 independent experiments, each with ≥4 biological replicates.

In complementary studies, the MrgprD agonist β-alanine (Guo Changxiong et al., 2023) was injected intradermally in the ear for pharmacological NP1 neuron activation before topical application of Vaseline or 5% IMQ cream (**Fig. S2A**). Consistent with our optogenetic approach, stimulation of MrgprD+ afferents using β-alanine also significantly attenuated the IMQ-driven increase in ear thickness (**Fig. S2B**). No increase in ear thickness in vehicle- or β-alanine-treated mice exposed to Vaseline was observed (**Fig. S2B**). Thus, optogenetic or pharmacologic activation of NP1 neurons significantly reduced skin inflammation caused by IMQ.

To test whether increased ear thickness reflected a local increase in epidermal thickness and skin pathology, we performed H&E staining of formalin-fixed paraffin-embedded (FFPE) skin biopsies collected after 5 days of Vaseline or IMQ exposure in control or MrgprD-ChR2 strains. While IMQ treatment substantially increased the epidermal thickness of both control and MrgprD-ChR2 groups compared to their respective Vaseline controls, IMQ-treated MrgprD-ChR2 mice had significantly less epidermal thickness compared to IMQ-exposed controls (**Fig. 4C,D**). Clinical scoring of these histological sections revealed lower hyperkeratosis, subcorneal pustules, acanthosis, basal hyperplasia, and neutrophil and mast cell infiltrates in IMQ-treated MrgprD-ChR2 mice compared to controls (**Fig. 4E**). Similarly, β-alanine treatment also reduced the epidermal thickening induced by IMQ compared to vehicle-treated controls (**Fig. S2C,D**). Flow cytometry analysis of skin cell suspensions confirmed that the IMQ-induced increase in skin neutrophils was significantly reduced in both frequency and number when MrgprD-ChR2 or β-alanine-treated mice were compared to their respective controls (**Fig. 4F,G; Fig. S3A**).

### IL-17 cytokine responses induced by IMQ are reduced by stimulating NP1 neurons

The ability of local NP1 neuron stimulation to reduce IMQ-induced skin pathology prompted us to investigate whether inflammatory cell subsets and/or cytokine secretion patterns were also altered. NP1 activation using either approach significantly blunted IL-17A, IL-17F, and IL-22 supernatant levels from sdLNs restimulated with αCD3/αCD28 (**Fig. 5A, Fig. S3B**). Notably, intracellular cytokine staining to identify IL-17+ cells within sdLNs at 5 days of IMQ application revealed that although IMQ-treated control mice had a nearly 6-fold increase in the percentage of total IL-17+ cells compared to Vaseline controls, there was only a 2-fold increase in IMQ-exposed MrgprD-ChR2 mice (**Fig. 5B,C**). Although γδ T cells represented the majority of IL-17-expressing cells in control and MrgprD-ChR2 strains treated with IMQ compared to Vaseline controls, the proportion of IL-17+ γδ T cells within IMQ-treated MrgprD-ChR2 mice was reduced compared to controls (**Fig. 5D,E**). CD4+ T helper (Th) cells and innate lymphoid cells (ILCs) comprised moderate proportions of IL-17+ cells, with CD8+ T cells and Foxp3+ T regulatory cells expressing low levels of IL-17. Activation of NP1 neurons with optogenetics or β-alanine significantly blunted the overall IMQ-driven accumulation of γδ T cells in both percentage and number within sdLNs (**Fig. 5F, Fig. S3C**).

**Figure 5.**
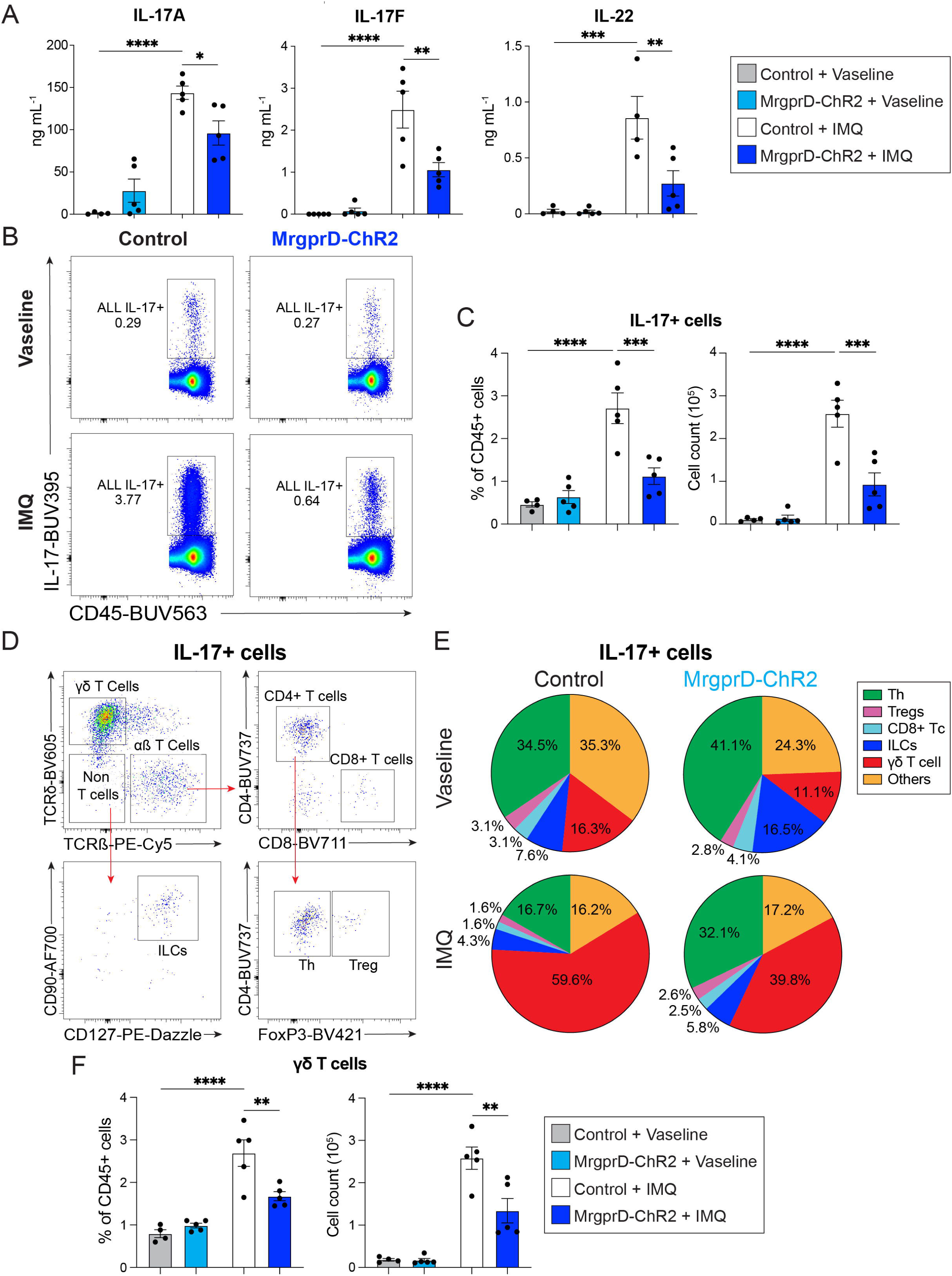
NP1 neurons suppress skin IL-17 cytokine responses induced by IMQ. (**A**), Concentration of IL-17A, IL-17F, and IL-22 in cell-free supernatants from restimulated (αCD3/αCD28, 72hrs) sdLNs from control or MrgprD-ChR2 mice treated for 5 days with Vaseline or 5% IMQ. (**B,C**), Representative dot plots, percentage, and absolute numbers of IL-17+ cells from sdLNs restimulated with PMA/Ionomycin for 5 hours and analyzed by flow cytometry. (**D,E**), Flow cytometric gating strategy and pie charts of average percentages of cellular subsets within IL-17+ cells from B. (**F**), Percentage and absolute numbers of sdLNs γδ T cells (CD45+ CD90+ CD127+ TCRδ+ TCRβ-) from control or MrgprD-ChR2 mice treated for 5 days with Vaseline or 5% IMQ. Bar charts depict mean ± s.e.m (n=4-5). *P* values were determined by One-way ANOVA with Bonferroni post hoc correction. **P<0.01, ***P<0.001, ****P<0.0001. Representative of 2-3 independent experiments, each with ≥4 biological replicates.

### Stimulation of NP1 neurons alters the activation profile of γδ T cells

Although TRPV1+ and MrgprA3+ neurons can induce IL-17 secretion by γδ T cells (Inclan-Rico et al., 2024, Inclan-Rico et al., 2025, Kashem et al., 2015, Riol-Blanco et al., 2014), whether stimulation of MrgprD+ afferents impacts γδ T cell biology remains unknown. FlowSOM was used to evaluate cytokine/cytokine receptor, transcription factor, and proliferation profiles of γδ T cells in the context of NP1 afferent stimulation. Our analysis focused on expression levels of Ki67, IL-17, Areg, IFNγ, IL-13, GATA3, ST2, and TNF to distinguish between distinct γδ T cell phenotypes. Comparison of Vaseline-treated groups revealed moderate separation of γδ T cell clusters due to NP1 afferent stimulation in the MrgprD-ChR2 group, with slight, but notable reductions in Ki-67, IL-17, and Areg (**Fig. S4A-C**). Likewise, there was substantial overlap when comparing Vaseline-treated groups in the context of pharmacological NP1 stimulation with β-alanine (**Fig. S4D-F**). Further, FlowSOM analysis of sdLN γδ T cell populations revealed a clear separation between IMQ-treated control and MrgprD-ChR2 groups, with enrichment of IL-17, Areg, and Ki-67 populations in controls, but IL-13 and GATA3 enrichment in γδ T cells of MrgprD-ChR2 mice (**Fig. 6A-C**). Twelve clusters were identified, with Clusters 1-3 expressing high levels of IL-17, Ki-67, and Areg enriched in controls, whereas Clusters 9-12 were enriched in MrgprD-ChR2 mice showing moderate-to-high levels of IL-13 and GATA3 (**Fig. 6D-F**). Clusters 4-8 were similarly represented in control or MrgprD-ChR2 mice. Regarding β-alanine-induced NP1 activation, γδ T cells from IMQ-exposed mice injected with vehicle were enriched in IL-17, Areg, and Ki-67 expression, whereas γδ T cells in β-alanine-treated mice had reduced expression of these markers and moderately increased IL-13 expression levels (**Fig. S5A-C**). Of the nine clusters identified, Clusters 6-9 were elevated in response to β-alanine treatment, displaying low to moderate expression of IL-17, Areg, and Ki-67 (**Fig. S5D-F**). On the other hand, Clusters 1-3 that expressed high levels of these cytokines were increased in vehicle-treated mice (**Fig. S5D-F**). Collectively, these data indicated that activation of NP1 neurons through optogenetic or pharmacologic approaches shifts the phenotype of γδ T cells elicited during psoriasiform inflammation.

**Figure 6.**
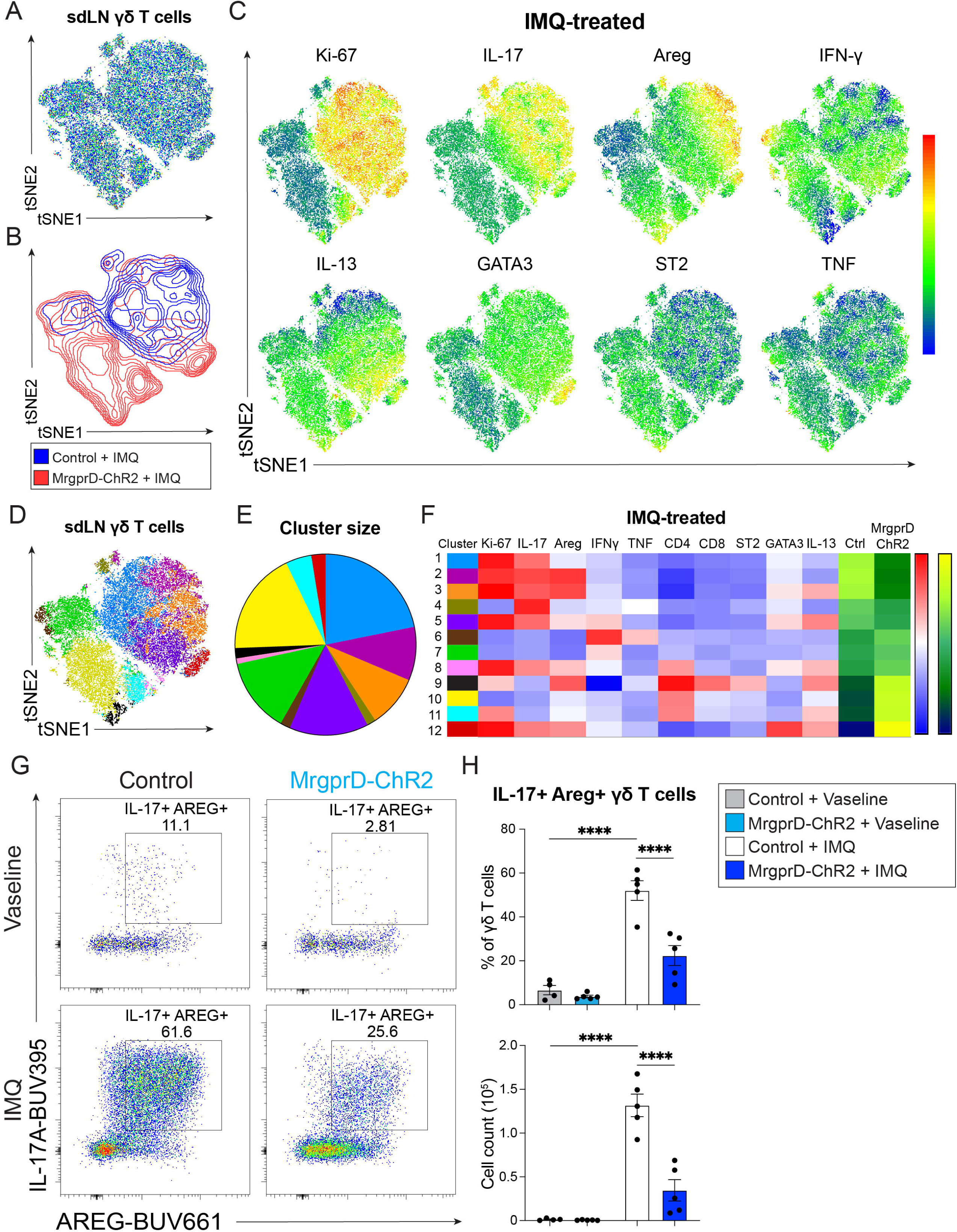
NP1 neurons modulate the activation profile of γδ T cells during psoriasiform dermatitis. (**A,B**), t-SNE dot plots and counterplots of concatenated γδ T cells from sdLNs of control or MrgprD-ChR2 mice treated for 5 days with 5% IMQ cream. (**C**), Expression of cytokines, ST2, GATA-3 and Ki67 in t-SNE dot plots from A. (**D-F**), FlowSOM analysis, cluster sizes, and heatmaps of γδ T cell clusters. (**G,H**), Representative dot plots, percentages, and absolute numbers (mean ± s.e.m (n=4-5)) of IL-17^+^ Areg^+^ or Ki-67^+^ γδ T cells quantified with traditional flow cytometric gating. *P* values were determined by One-way ANOVA with Bonferroni post hoc correction. ****P<0.001. Representative of 2-3 independent experiments, each with ≥3 biological replicates.

Traditional flow cytometric analysis confirmed that γδ T cells co-expressing IL-17 and Areg were significantly increased in IMQ-treated controls compared to Vaseline-treated (4-5-fold) and IMQ-treated MrgprD-ChR2 mice (2-3-fold) in both frequency and number (**Fig. 6G,H**). Similarly, the increased proportion and number of proliferating Ki67^+^ γδ T cells due to IMQ exposure was significantly reduced via optogenetic NP1 neuron activation (**Fig. S5G**). Activating NP1 neurons using optogenetics before IMQ exposure led to an increased frequency of IL-13^+^ γδ T cells (**Fig. S5H**). Although less pronounced, β-alanine treatment during IMQ application also reduced the levels of IL-17^+^Areg^+^ γδ T cells, whereas changes in Ki-67+ γδ T cells were moderate and did not reach statistical significance (**Fig. S6A,B**). However, β-alanine-induced activation of NP1 neurons significantly increased IL-13^+^ γδ T cells in both frequency and number (**Supp. Fig 6C**). Regarding γδ T cell subsets, Vγ4+ (50-60%) and Vγ6+(35-45%) accounted for most IL-17^+^Areg^+^ or IL-13^+^ γδ T cells (**Fig S6D**). Lastly, using ELISA to directly measure the impact of NP1 activation on IL-13 secretion from sdLNs revealed significantly higher levels in cultures from MrgprD-ChR2 mice as compared to controls, irrespective of Vaseline or IMQ treatment (**Fig. S6E**). β alanine treatment also significantly increased IL-13 secretion in the context of IMQ exposure (**Fig. S6F**). Collectively, these data show that activation of NP1 neurons increased Type 2 cytokine secretion and reduced IMQ-induced IL-17 cytokine production.

### NP1 neuron activation induces IL-10 that is necessary to suppress psoriasiform inflammation

The immunoregulatory cytokine IL-10 has been shown to suppress IMQ-induced dermatopathology (Jin et al., 2018). Given that activation of MrgprD+ neurons suppressed psoriasiform immunopathology, we postulated a potential role for IL-10. Measurement of IL-10 levels in supernatants from restimulated sdLNs revealed that optogenetic stimulation of MrgprD+ neurons increased IL-10 levels in both Vaseline- and IMQ-exposed mice (**Fig. 7A**). Similarly, β−alanine treatment significantly increased IL-10 secretion from sdLNs during IMQ application (**Fig. 7B**). To formally test whether IL-10R signaling was necessary for the NP1-induced suppression of IMQ-evoked dermatitis, WT mice were treated with Isotype or neutralizing IL-10RmAb at three days and one day before IMQ application and β−alanine intradermal treatment (**Fig. 7C**). While β−alanine treatment significantly attenuated IMQ-induced ear swelling in isotype-treated controls, administration of anti-IL-10R antibody and β−alanine resulted in equivalent ear thickness as vehicle-treated controls **(Fig. 7D**). Comparably, H&E staining of FFPE ear sections showed that β−alanine treatment reduced IMQ-induced epidermal hyperplasia, but not upon co-administration of anti-IL-10R mAb (**Fig. 7E,F**). Importantly, anti-IL-10R mAb treatment reversed the ability of β−alanine to suppress the accumulation of IL-17^+^ Areg^+^ and Ki-67^+^ γδ T cells normally induced by IMQ exposure (**Fig. 7G-I**). Collectively, these data indicated that MrgprD+ neurons blocked IMQ-induced dermatitis, at least in part, through an IL-10-dependent mechanism.

**Figure 7.**
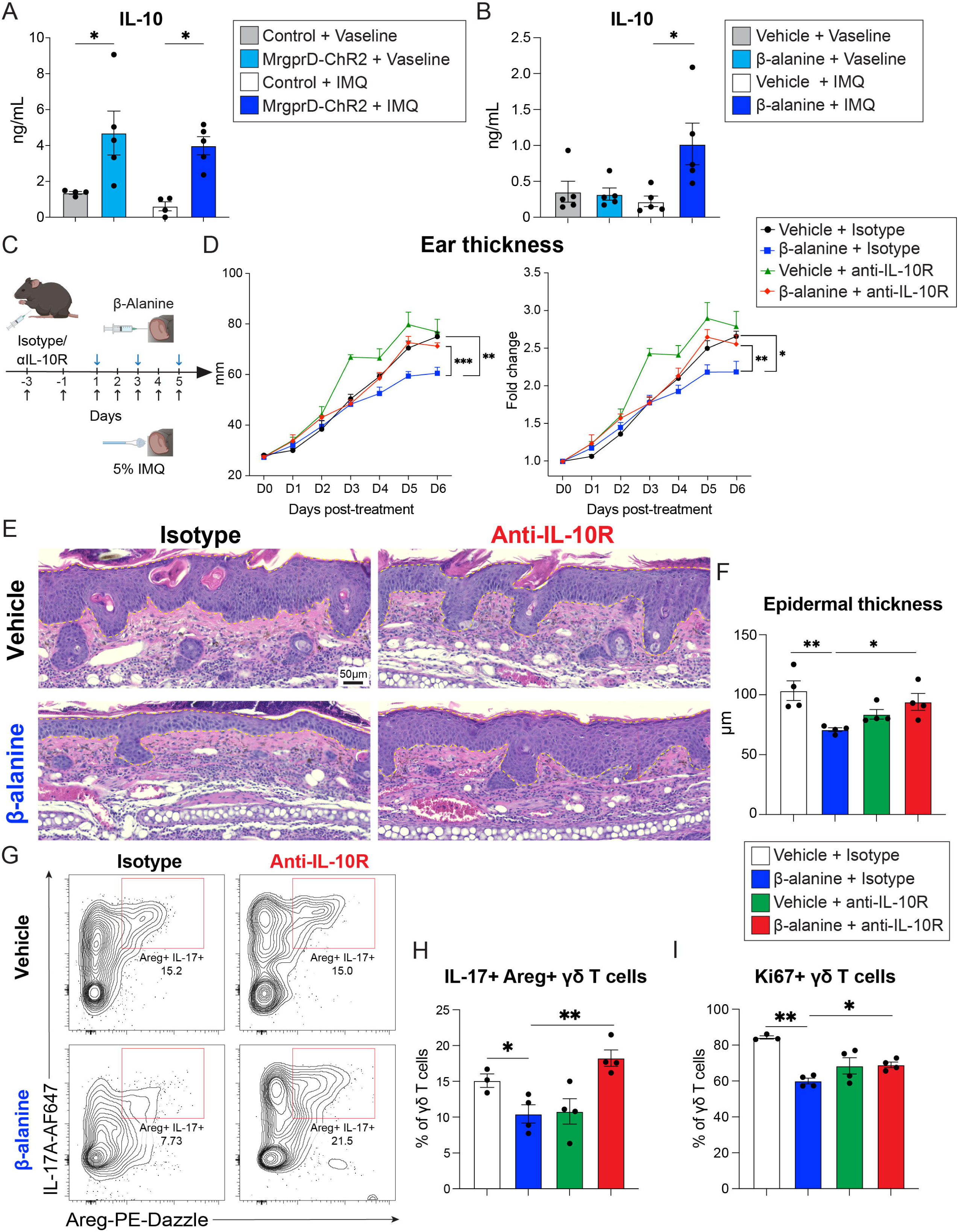
IL-10 is required for the suppression of IMQ-induced skin inflammation evoked by NP1 neurons. (**A**), Concentration of IL-10 in cell-free supernatants from re-stimulated (αCD3/αCD28, 72hrs) sdLNs from control and MrgprD-ChR2 mice or (**B**), vehicle- and β-alanine-treated mice after 5 days of Vaseline or 5% IMQ application. (**C**), Mice were injected with isotype control or anti-IL-10R neutralizing antibody 3 and 1 day before vehicle- and β-alanine-treatment and IMQ application. (**D**), Raw values and fold change of ear thickness measured daily. (**E,F**), H&E staining and epidermal thickness quantification of FFPE ear skin sections after 5 days of IMQ exposure. Dotted lines indicate epidermal thickness. (**G-I**), Representative dot plots and percentages of IL-17^+^ Areg^+^ or Ki-67^+^ γδ T cells from sdLNs collected 5 days after IMQ. Line or bar graphs depict mean ± s.e.m (n=3-5). *P* values were determined by One-way ANOVA with Bonferroni post hoc correction, or Two-way ANOVA with Tukey’s for post hoc test. *P<0.05, **P<0.01, ***P<0.001. Representative of 2-3 independent experiments, each with ≥4 biological replicates.

### Stimulation of NP2 neurons exacerbates IMQ-induced inflammation

NP neuron subsets likely serve distinct biological roles given the distinct cadre of neuropeptides and neurotransmitters that they produce. For instance, NP1 afferents transmit β−alanine-induced itch, mechanical pain, or scratch suppression of itch; whereas NP2 neurons mostly transmit CQ-evoked pruritus (Cranfill et al., 2025, Guo Changxiong et al., 2023, Han et al., 2013, Liu et al., 2012). We postulated that NP2 neurons could potentially serve a disparate role in IMQ-induced dermatitis as compared to MrgprD+ neurons. Using a MrgprA3^Cre^: Ai9-Rosa26^tdTomato^ line for visualization of MrgprA3 skin afferents, data show that MrgprA3+ afferents were less frequent in the epidermis and mostly localized in the dermis (**Fig. 8A**). Unlike NP1 afferents, the density of MrgprA3 innervation was not significantly different between Vaseline-and IMQ-exposed groups (**Fig. 8A,B**). The biological role of MrgprA3 afferents during IMQ-induced cutaneous inflammation was evaluated in photostimulated control or MrgprA3-ChR2 mice (**Fig. 8C**). Daily photostimulation of MrgprA3-ChR2 groups increased IMQ-driven ear swelling compared to controls (**Fig. 8D**). Consistently, H&E staining of ear sections showed that photostimulated MrgprA3-ChR2 mice had increased IMQ- driven ear thickness relative to controls (**Fig. 8E,F**). MrgprA3 activation also augmented IL-17 cytokine responses, shown by flow cytometric and t-SNE analysis of γδ T cells in sdLNs, revealing that control and MrgprA3-ChR2 groups had distinct cell clusters (**Fig. 8G**). γδ T cell clusters in MrgprA3-ChR2 mice were enriched for IL-17, Areg, and Ki67+ cells, whereas cells in controls had higher IFN-γ and TNF expression (**Fig. 8H-K**). Traditional flow cytometric analysis confirmed that optogenetic stimulation of MrgprA3 afferents increased IL-17+ Areg+ and Ki67+ γδ T cell percentages and numbers during IMQ exposure over controls (**Fig. 8L,M**). Consistently, IL-17A and IL-22 supernatant levels of restimulated sdLNs from IMQ-treated MrgprA3-ChR2 mice were significantly higher than those in controls (**Fig. 8N**). Taken together, these data indicate that NP1 and NP2 skin afferents exert opposing effects on IMQ-driven cutaneous immunopathology and IL-17 responses.

**Figure 8.**
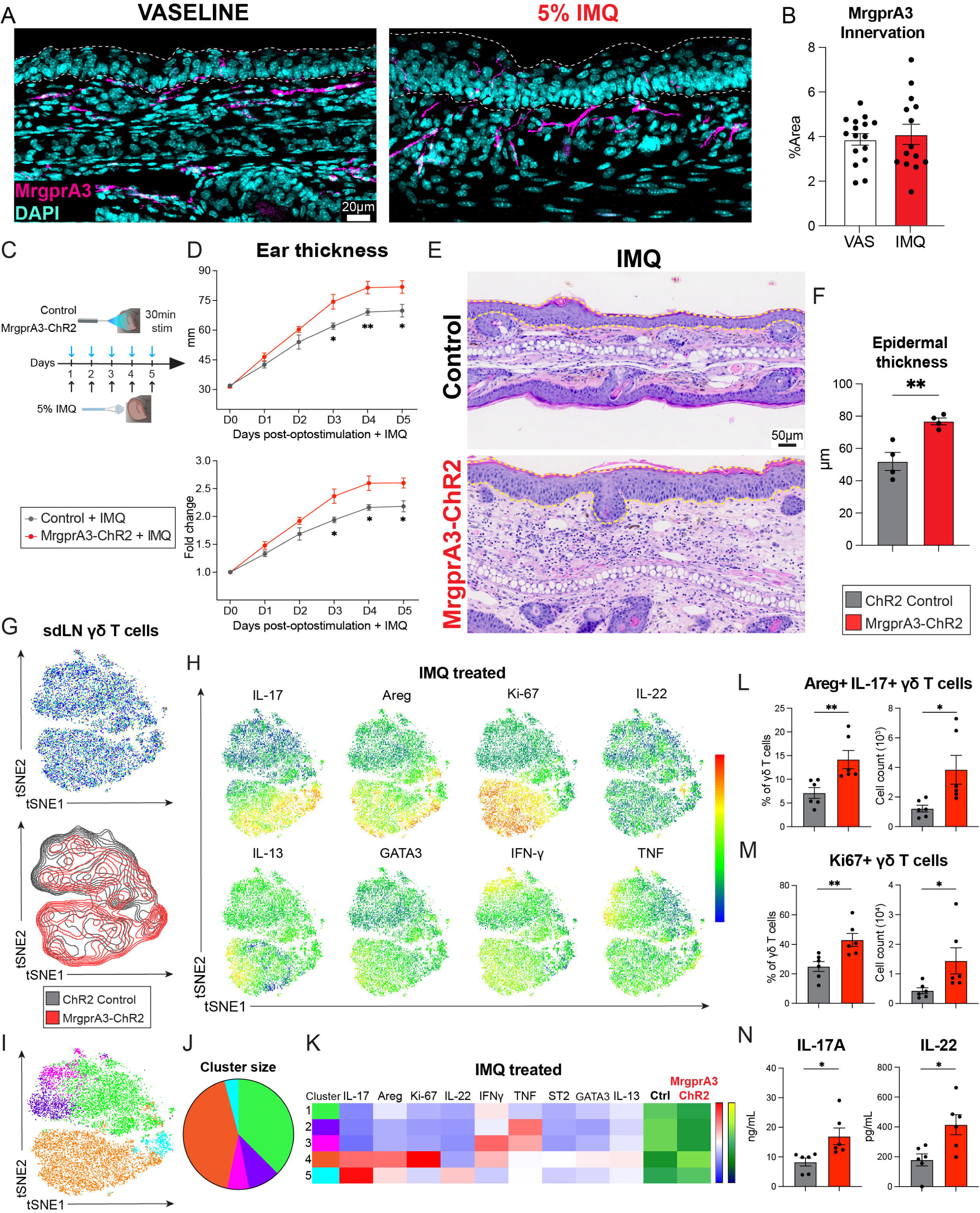
Optogenetic stimulation of MrgprA3+ afferents worsens psoriasiform inflammation. (**A**), IFA staining of MrgprA3-tdTomato+ afferents in ear skin sections treated with Vaseline or 5% IMQ cream for 5 consecutive days. (**B**), Mean ± s.e.m (n=14-16) of the percentage Area covered by MrgprA3-tdTomato+ nerves. (**C**), Experimental approach for optogenetic ear stimulation of ChR2 control or MrgprA3-ChR2 mice followed by topical application of 5% IMQ cream. (**D**), Raw values and fold change of ear thickness measured daily. (**E,F**), H&E staining and epidermal thickness quantification of FFPE ear skin sections from photostimulated control or MrgprA3-ChR2 mice after 5 days of IMQ treatment. Dotted lines indicate epidermal thickness. (**G**), t-SNE dot plots and counterplots of concatenated γδ T cells from sdLNs of control or MrgprA3-ChR2 mice after IMQ treatment. (**H**), Expression of cytokines, Ki67, and GATA3 in t-SNE dot plots from G. (**I-K**), FlowSOM analysis, cluster sizes, and heatmaps of γδ T cell clusters from G. (**L,M**), Percentage and absolute numbers of IL-17^+^ Areg^+^ or Ki-67^+^ γδ T cells quantified with traditional flow cytometric gating. (**N**), Concentration of IL-17A, and IL-22 in cell-free supernatants from skin-draining lymph nodes (sdLNs) stimulated with αCD3/αCD28 antibodies for 72hrs. Bar graphs depict mean ± s.e.m (n=4-5). *P* values were determined by two-tailed Student’s t-tests or Two-way ANOVA with Tukey’s for post hoc test. *P<0.05, **P<0.01. Representative of 2-3 independent experiments, each with ≥4 biological replicates.

## Discussion

Psoriasis is a prevalent autoimmune disorder that manifests with intense pruritus and skin γδT17-associated inflammation, but the contribution(s) of discrete sensory neuron subsets to disease pathology remained unclear. This study shows that human itch-transmitting NP1 and NP2 neuron subsets are enriched in PRR and cytokine receptor expression associated with psoriasis at baseline, suggesting that NP neurons may be engaged during psoriasis through signaling through one or more of these receptors. In support of this, *in silico* analysis showed profound transcriptional alterations of neuropeptides, neurotransmitters, and their receptors expressed in cutaneous lymphocytes in two publicly available datasets from human patients with active psoriatic lesions, as well as in a mouse model that resembles human psoriasis. Pain-inducing Nav1.8+ and TRPV1+ neurons were shown to promote IL-17 responses in the IMQ-driven murine model of psoriasiform dermatitis (Hanč et al., Riol-Blanco et al., 2014), but it had not been considered whether other types of sensory afferents lacking TRPV1 expression could have divergent role(s) in this disease. MrgprD+ (NP1) afferents densely innervate the epidermis at baseline but were lost within 5 days of IMQ administration, whereas MrgprA3+ afferents were less frequent in the epidermis and innervation was not altered after IMQ. Unexpectedly, NP1 neurons served an immunoregulatory role in IMQ-induced inflammation, shown by the exacerbated epidermal thickness, psoriatic clinical scores, and γδT17-responses following NP1 ablation. Conversely, two gain-of-function strategies using preemptive NP1 activation with optogenetics or β−alanine markedly blunted IMQ-driven skin pathology and significantly reduced γδ T cell populations that co-expressed psoriasis-driving cytokines (IL-17 and Areg). Instead, NP1 activation caused expansion of IL-13+ γδ T cells and IL-10 secretion, with IL-10R signaling serving a key role in the inhibition of psoriasiform inflammation. On the other hand, optogenetic stimulation of NP2 (MrgprA3+) afferents exacerbated IMQ-induced dermatitis and γδT17 cell responses. Altogether, this highlights how molecularly distinct skin sensory neuron subsets (i.e., NP1 vs. NP2) can differentially regulate cutaneous immunopathology and γδ T cell biology.

Although psoriasis and psoriatic arthritis were recently suggested to increase the risk of polyneuropathy (Doneddu et al., 2024), conflicting reports have shown increased or decreased skin innervation in psoriatic lesions(Chen et al., 2022, El-Nour et al., 2009, Kim et al., 2014, Pergolizzi et al., 1998, Taneda et al., 2011). MrgprD+ afferents constitute 60% of free nerve endings in the epidermis, and their innervation projects into the stratum granulosum, only 10μm away from the most external layer of the epidermis, the stratum corneum (Zylka et al., 2005). Five days of IMQ exposure were sufficient to reduce epidermal innervation as well as the density of MrgprD+ afferents, although the remaining fibers still extended through thickened epidermal layers. These findings suggest that NP1 neurons are specifically reduced during psoriasiform dermatitis, but they retain their ability to extend through newly formed epidermal layers. The increased psoriasiform pathology and inflammation detected in animals lacking MrgprD+ neurons suggests that NP1 afferents tonically restrict γδT17 cells in the skin. These results are consistent with reduced *Mrgprd* expression in the DRG after IMQ treatment, correlated with a substantial increase in spontaneous itching (Sakai et al., 2016). Interestingly, reduced MrgprD+ neurons and increased itching were also reported in the context of an SABDE-induced contact dermatitis mouse model that lacked *Trpc3* expression (Beattie et al., 2022). In this study, NP1 neuron loss was proposed to be due to increased sensitivity to neuronal excitotoxicity in the absence of Trpc3. More recently, we showed that MrgprD+ NP1 afferents are required for sensing scratching-induced mechanical stimuli to suppress further scratching (Cranfill, 2025). NP1 afferent ablation resulted in augmented scratching in response to chloroquine-induced itch, sensed by NP2 afferents. Although we did not formally quantify scratching behavior in this study, we hypothesize that reduction of NP1 afferents in psoriasiform dermatitis may lead to increased scratching, which could further augment skin immunopathology. A comprehensive evaluation of the density of specific NP afferents in the skin over the course of disease progression in psoriatic patients is warranted to understand how local inflammation impacts innervation by discrete subsets of sensory neurons.

It is known that pro-inflammatory cytokines such as TNF, IL-1β, and IL-6 can cause nociceptor sensitization and potentially epidermal nerve loss (Binshtok et al., 2008, Constantin et al., 2008, Oprée and Kress, 2000). We postulate that inflammatory mediators released during psoriasis may signal through specific receptors expressed in NP1 and NP2 skin-innervating afferents and potentially alter their expression and/or function regarding neuropeptide/neurotransmitters. Thus, we analyzed the expression of neuropeptides, neurotransmitters, and their receptors in publicly available RNA-seq data from healthy skin, non-lesional biopsies, and active psoriatic lesions. This analysis revealed profound upregulation of CGRP, NMB, and somatostatin in active psoriatic lesions, which are expressed by NP and peptidergic neuron subsets of both humans and mice (Guo Changxiong et al., 2023, McCoy et al., 2013, Xing et al., 2020, Yu et al., 2024). On the other hand, Substance P, VIP, NMU, and tyrosine hydroxylase, which controls serotonin synthesis, were downregulated in psoriatic biopsies. Lesions of psoriatic lesions also had increased expression of NMU, CGRP, and Substance P receptors, whereas receptors for VIP, Neuropeptide FF, and glutamate were curiously downregulated. Analysis of our previously published deep RNA-seq data of human DRG neurons, focusing on itch-transmitting NP neurons, revealed that among NP subsets, NP1 neurons were most enriched in PRRs, including the receptor for IMQ, TLR7; suggesting that NP1 afferents could potentially respond to IMQ applied to the skin surface. Although human NP2 neurons expressed fewer PRRs, they notably were enriched in the cytosolic receptor MDA5, which is associated with psoriatic disease progression (Coto-Segura et al., 2023). NP1 neurons were also enriched in receptors for IL-1β, IL-6, IL-17A/F, IL-23, TNF, and type I/II IFNs, which are all recognized to promote psoriasis pathology (Armstrong et al., 2025), whereas NP2 afferents expressed more discrete levels of IL-1β, IL-6, and TNF receptors at baseline. The specific role for each of these PRR and cytokine signaling pathways in NP1 and NP2 biology will require development of precise targeting of individual pathways using subset-specific genetic tools.

γδ T cells with a morphology resembling dendritic epidermal T cells (DETC) localized closely to NP1 afferents in the epidermis at baseline, but IMQ exposure increased the distance between them and induced their ‘rounding’ associated with an activated phenotype (Strid et al., 2011). Despite MHC-II^+^ myeloid APCs being mostly located in the dermis, their distance with from NP1 neurons increased after IMQ treatment. We and others have previously identified the proximity of hematopoietic cells, such as APCs and T cells, to sensory neurons in both the dermis and epidermis (Deng et al., 2023, Enamorado et al., 2023, Fassett et al., Inclan-Rico et al., 2024, Inclan-Rico et al., 2025, Liu et al., 2025, Zhang et al., 2021). Thus, NP1 neurons may regulate the functions of both γδ T cell subsets and myeloid APCs at homeostasis, but the loss of NP1 afferents in IMQ-exposed epidermis results in the loss of such regulation. Epidermal γδ T cells can also regulate the functions of itch-inducing peptidergic neurons via their secretion of IL-3 (Flayer et al., 2024), suggesting that bidirectional communication between immune cells and peripheral nerves regulates skin inflammation. Dermal γδ T cells, but not DETCs, that express Vγ4 and Vγ6 are well established as the main source of IL-17 that drives skin pathology induced by IMQ (Cai et al., 2011, Cai et al., 2014, Gray et al., 2013, Pantelyushin et al., 2012, Zhu et al., 2017). Nonetheless, Vγ4 γδ T cells have a greater capacity to secrete IL-17 and are proposed to migrate into the skin from the lymph node, where they drive the skin remodeling induced by IMQ (Gray et al., 2013, Ramírez-Valle et al., 2015). Multiparametric flow cytometry analysis that focused on sdLN γδ T cells revealed that γδ T cells are heterogeneous, with low baseline levels of IL-17 and Ki-67, with IMQ inducing robust increases in these parameters. Notably, subsets of γδ T cells from IMQ-treated mice co-expressed IL-17 and Areg, both of which are known to promote keratinocyte proliferation, maturation, and chemokine secretion (Cook et al., 1997, Jia et al., 2018, Rizzo et al., 2011). Areg+ IL-17+ γδ T cells were previously shown to promote wound healing and maintain the integrity of the oral mucosa (Krishnan et al., 2018). While Areg secretion by keratinocytes is well known to drive psoriasis (Cook et al., 1997), whether IMQ-induced Areg expression by γδ T cells serves a distinctly pathogenic role requires further investigation.

One of the most remarkable findings of this study is the robust effect of NP1 neuron stimulation on the activation profile of γδ T cells. NP1 neurons not only suppressed IL-17, Areg, and Ki-67 in γδ T cells, but they also increased their expression of IL-13 and GATA3, suggesting that NP1 neurons may reprogram γδ T cells to suppress IMQ-induced inflammation. Consistent with the studies cited above, IL-17^+^ Areg^+^ γδ T cells expressed Vγ4 or Vγ6 in similar proportions, which was also observed in IL-13^+^ γδ T cells, suggesting that NP1 stimulation reprograms similar γδ T cell subsets rather than recruiting different cell lineages. GATA3 expression was previously reported in tissue-resident γδ T cells from neonatal pigs, although its functional role was unclear since it did not induce IL-4 secretion by γδ T cells (Rodríguez-Gómez et al., 2019). Although our study revealed IL-13 induction in sdLN γδ T cells, TCR signaling in DETCs was previously reported to induce IL-13 secretion, which was proposed to maintain epithelial barrier integrity and promote epithelial repair (Dalessandri et al., 2016, Ibusuki et al., 2024). IL-13-expressing DETCs have been proposed to mediate the early stress-surveillance response in the epidermis through communication via the receptor NKG2D (Strid et al., 2011). Epidemiologic data have shown that use of IL-13 inhibitors for atopic dermatitis was previously shown to induce psoriatic episodes (Grolleau et al., 2023, Zhao et al., 2024), suggesting that promoting IL-13 responses may reduce psoriatic pathology. In perspective, NP1 neuron stimulation may reprogram the functions of γδ T cells, driving them away from γδT17 and Areg+ subsets during IMQ-induced dermatitis and potentially restoring a skin immunosurveillance phenotype (McKenzie et al., 2022, Sumaria et al., 2011) characterized by IL-13 and GATA3 expression. However, whether IL-13+ γδ T cells in the sdLN induced upon NP1 neuron stimulation migrate into the skin or act remotely through their cytokine secretion warrants further clarification. On the other hand, NP1 neurons could potentially regulate skin γδ T cells directly, however, recirculation of γδ T cells from the skin into sdLNs has only been reported in bovine and ovine systems, so whether this process occurs in murine skin is unclear (Geherin et al., 2013, Ugur et al., 2018, Van Rhijn et al., 2007, Vrieling et al., 2012).

Increased levels of several pruritic mediators, including IL-31, oncostatin M, histamine, and serotonin, were previously reported in human psoriatic lesions or in the mouse model of IMQ-induced dermatitis (Hashimoto et al., 2019, Krogstad et al., 1997, Petersen et al., 1998, Sakai et al., 2016, Tseng and Hoon). This suggests that psoriatic itch is likely mediated by several sensory afferent subsets. In support of this, the neuropeptides CGRP and Substance P are increased in serum from psoriatic patients (Narbutt et al., 2013, Reich et al., 2007), and afferents expressing Nav1.8 or TRPV1, including NP and peptidergic neurons, were previously shown to promote IMQ-induced dermatitis (Kim et al., 2011, Riol-Blanco et al., 2014, Zhou et al., 2018). In order to avoid scratch-associated effects that could confound our interpretations, our optogenetic experiments were always performed in anesthetized mice (4% Isofluorane), which prevented scratching during the photostimulation session. Scratching was not quantified after photostimulation, although optogenetic stimulation of MrgprD+ afferents in the paw of awake mice was shown to induce avoidance behavior or drive mechanical allodynia (Olson et al., 2017, Wang et al., 2023); thus, it is unclear if our photostimulation of MrgprD+ afferents in the ear of awake mice evokes scratching. Beyond their capacity to transmit β-alanine-induced itch, polymodal NP1 neurons transmit different sensory stimuli such as mechanical pain, nerve-injury mechanical allodynia, and scratch sensing to negatively regulate itch-induced behavior(Cranfill et al., 2025, Guo Changxiong et al., 2023, Wang et al., 2023). This differential activity is determined at least in part by the mediators released by NP1 afferents, as shown by studies showing that glutamate was required to evoke mechanical pain and itch, whereas NMB was exclusively necessary for itch transmission (Guo Changxiong et al., 2023). NP1 neurons were previously shown to suppress mast cell responses during cutaneous bacterial infections through their glutamate release (Zhang et al., 2021). Interestingly, our transcriptional analysis of skin lymphocytes during a mouse model where keratinocyte-specific *Ube2l3* deletion leads to psoriasis-like pathology (Chen et al., 2025), γδT17 cells were enriched for NMBR and several genes for glutamate receptor signaling such as *Grin1, Grin2c*, and *Grin3b*. γδT17 cells also expressed high levels of bradykinin and endogenous opioid receptors, but the diversity of mediators being released by NP1 afferents during psoriatic inflammation remains unclear. We cannot rule of the possibility that NP1 neurons also regulate myeloid APCs and their secretion of IL-23 and IL-1β, which are well known to drive IL-17 γδ T cell secretion. Thus, it remains unclear whether NP1 neurons regulate the functions of γδ T cells directly or indirectly through modulating APCs. We recently demonstrated that myeloid APC expressing the alarmin cytokine IL-33 have a significant role in driving host protective γδT cell responses against skin-penetrating helminths (Jean et al., 2025)

Most notably, NP1 neuron stimulation promoted IL-10 secretion in IMQ-exposed mice, and this IL-10 was essential for the ameliorative effect of NP1 stimulation on psoriasiform immunopathology. While rare, IL-10-producing γδ T cells were shown to suppress TNF production in CD8+ T cells and protect liver function during *Listeria* infection (Rhodes et al., 2008). Further, a unique population of circulating Vγ9^-^ γδ T cells enriched in IL-10 expression was previously found in patients with ‘naturally acquired immunity’ against malaria (Taniguchi et al., 2017). While γδ T cells may potentially be a source of IL-10, B cells that produce IL-10 inversely correlate with the severity of human psoriasis, and IL-10 has proven successful as an anti-psoriatic treatment (Asadullah et al., 1999, Mavropoulos et al., 2017, Reich et al., 1998). Accordingly, B cell deficiency exacerbated IMQ-induced inflammation, which was dampened after transfer of regulatory B cells that express IL-10, negatively regulated by the transcription factor Nfatc1 (Alrefai et al., 2016, Yanaba et al., 2013). Similarly, several approaches that ameliorate psoriasiform dermatitis report increases in Foxp3+ Tregs and IL-10 (Hau et al., 2018, Nakao et al., 2020, Schwarz et al., 2021). Further studies will define the cellular sources and molecular mechanism of IL-10 induction by NP1 neurons. Of note, our approaches to stimulate NP1 neurons were performed simultaneously with the induction of skin pathology, as this model requires continuous IMQ application. Thus, the therapeutic potential of NP1 neuron stimulation in preclinical models of established psoriatic pathology or in chronic stages of human disease remains to be determined.

Lastly, optogenetic stimulation of MrgprA3+ NP2 afferents exacerbated IMQ-induced skin immunopathology, which is consistent with our previous work showing that MrgprA3+ neurons are sufficient to induce skin γδT17 cells (Inclan-Rico et al., 2024). Subsets of MrgprA3 neurons express the ion channel TRPV1 (Han et al., 2013, Solinski et al., 2019), which has also been shown to induce cutaneous IL-17-mediated γδ T cell responses upon optogenetic stimulation, although TRPV1 is also expressed in multiple afferent subsets like NP3 and PEP1 (Cohen et al., 2019, Inclan-Rico et al., 2025, Usoskin et al., 2015, Yu et al., 2024). The opposing effects of stimulating NP1 and NP2 neurons is likely due to their distinct molecular signature, which not only distinguishes the mediators they release, but also in their expression of PRR and cytokine receptors. In sum, our results propose a model in which NP1 and NP2 skin-innervating afferents are distinctly affected during psoriasiform dermatitis and opposingly regulate cutaneous immunopathology and IL-17-dominant responses induced by IMQ. This work highlights that molecularly distinct subsets of sensory neurons can have functionally disparate roles, which warrants further investigation into which neuron-derived molecules shape γδ T cell responses during cutaneous inflammation.

## Acknowledgments & Funding.

Diagrams created in BioRender. Inclan-Rico, J. (2025) https://BioRender.com/j03hyzp. J.M.I.R. is supported by the National Institutes of Health (1K99-AI187730-01A1), the Life Sciences Research Foundation, and the Skin Biology and Disease Research Core Pilot and Feasibility Grant of the University of Pennsylvania. W. L. is supported by National Institutes of Health (U19-NS135528, R01-NS131209, U01-EY034681 and R21-NS142656). D.R.H is supported by the National Institutes of Health (grant nos. T32-AI007532-24, R01-AI164715-01, U01-AI163062-01, R01-AI123173-05).

## Author contributions

Conceptualization: JMIR, DRH

Methodology: JMIR, CN, LYH, AS, HLR, UF, FM, HY

Investigation: JMIR, CN, LYH, AS, HLR, UF, FM, HY

Visualization: JMIR, DRH

Funding acquisition: JMIR, DRH

Supervision: WL, DRH

Writing – original draft: JMIR

Writing – review & editing: JMIR, WL, DRH

## Lead contact

Further information and requests for reagents may be directed to and will be fulfilled by the lead contacts Juan M. Inclan Rico and De’Broski R. Herbert.

## Data, Code, and Materials availability

All reagents generated or used in this study are available on request from the lead contact with a completed Materials Transfer Agreement. Information on reagents used in this study is available in the key resources **Table 1**.

## Declaration of Generative Artificial Intelligence (AI) or Large Language Models (LLMs)

No AI or LLM tools were used to prepare this manuscript.

## References

Alrefai H, Muhammad K, Rudolf R, Pham DAT, Klein-Hessling S, Patra AK, et al. NFATc1 supports imiquimod-induced skin inflammation by suppressing IL-10 synthesis in B cells. Nature Communications 2016;7(1):11724.

Armstrong AW, Blauvelt A, Callis Duffin K, Huang Y-H, Savage LJ, Guo L, et al. Psoriasis. Nature Reviews Disease Primers 2025;11(1):45.

Armstrong AW, Mehta MD, Schupp CW, Gondo GC, Bell SJ, Griffiths CEM. Psoriasis Prevalence in Adults in the United States. JAMA Dermatology 2021;157(8):940–6.

Armstrong AW, Read C. Pathophysiology, Clinical Presentation, and Treatment of Psoriasis: A Review. JAMA 2020;323(19):1945–60.

Asadullah K, Döcke W-D, Ebeling M, Friedrich M, Belbe G, Audring H, et al. Interleukin 10 Treatment of Psoriasis: Clinical Results of a Phase 2 Trial. Archives of Dermatology 1999;135(2):187–92.

Beattie K, Jiang H, Gautam M, MacVittie MK, Miller B, Ma M, et al. TRPC3 Antagonizes Pruritus in a Mouse Contact Dermatitis Model. Journal of Investigative Dermatology 2022;142(4):1136–44.

Binshtok AM, Wang H, Zimmermann K, Amaya F, Vardeh D, Shi L, et al. Nociceptors Are Interleukin-1β Sensors. The Journal of Neuroscience 2008;28(52):14062–73.

Cai Y, Fleming C, Yan J. New insights of T cells in the pathogenesis of psoriasis. Cell Mol Immunol 2012;9(4):302–9.

Cai Y, Shen X, Ding C, Qi C, Li K, Li X, et al. Pivotal Role of Dermal IL-17-Producing γδ T Cells in Skin Inflammation. Immunity 2011;35(4):596–610.

Cai Y, Xue F, Fleming C, Yang J, Ding C, Ma Y, et al. Differential developmental requirement and peripheral regulation for dermal Vγ4 and Vγ6T17 cells in health and inflammation. Nature Communications 2014;5(1):3986.

Chen S-Q, Chen X-Y, Cui Y-Z, Yan B-X, Zhou Y, Wang Z-Y, et al. Cutaneous nerve fibers participate in the progression of psoriasis by linking epidermal keratinocytes and immunocytes. Cellular and Molecular Life Sciences 2022;79(5):267.

Chen X-Y, Ye L-R, Fu N-C, Chen S-Q, Yan B-X, Zheng Y-X, et al. Single cell transcriptomics of human psoriasis and epidermal specific Ube2l3 deficient mice highlight CXCL16/CXCR6 involvement in psoriasis development. Nature Communications 2025;16(1):9084.

Cohen JA, Edwards TN, Liu AW, Hirai T, Jones MR, Wu J, et al. Cutaneous TRPV1^+^ Neurons Trigger Protective Innate Type 17 Anticipatory Immunity. Cell 2019;178(4):919–32.e14.

Constantin CE, Mair N, Sailer CA, Andratsch M, Xu Z-Z, Blumer MJF, et al. Endogenous Tumor Necrosis Factor α (TNFα) Requires TNF Receptor Type 2 to Generate Heat Hyperalgesia in a Mouse Cancer Model. The Journal of Neuroscience 2008;28(19):5072–81.

Cook PW, Piepkorn M, Clegg CH, Plowman GD, DeMay JM, Brown JR, et al. Transgenic expression of the human amphiregulin gene induces a psoriasis-like phenotype. The Journal of Clinical Investigation 1997;100(9):2286–94.

Coto-Segura P, Vázquez-Coto D, Velázquez-Cuervo L, García-Lago C, Coto E, Queiro R. The IFIH1/MDA5 rs1990760 Gene Variant (946Thr) Differentiates Early- vs. Late-Onset Skin Disease and Increases the Risk of Arthritis in a Spanish Cohort of Psoriasis. International journal of molecular sciences2023. p. 14803.

Cranfill SL, Yu H, Wang Y, Inclan-Rico JM, Janke E, Lezgiyeva K, et al. Asymmetric lateral habenula function and peripheral neural mechanisms in regulating itch-evoked scratching. Current Biology 2025;35(24):6180–90.e6.

Cranfill SY, Huasheng; Wang, Yingqi; Inclan-Rico, Juan M.; Janke, Emma; Lezgiyeva, Karina; Liu, Shibo; Chang, Annabel; Gooden, Steven; Baker, Jane; Shirvan, Sepenta; Wu, Qinxue; Bhattarai, Janardhan P.; Hill, Rose Z.; Ma, Minghong and Luo, Wenqin. Asymmetric Lateral Habenula Function and Peripheral Neural Mechanisms in Regulating Itch-Evoked Scratching. Current Biology 2025.

Dalessandri T, Crawford G, Hayes M, Castro Seoane R, Strid J. IL-13 from intraepithelial lymphocytes regulates tissue homeostasis and protects against carcinogenesis in the skin. Nature Communications 2016;7(1):12080.

de Koning HD, Rodijk-Olthuis D, van Vlijmen-Willems IMJJ, Joosten LAB, Netea MG, Schalkwijk J, et al. A Comprehensive Analysis of Pattern Recognition Receptors in Normal and Inflamed Human Epidermis: Upregulation of Dectin-1 in Psoriasis. Journal of Investigative Dermatology 2010;130(11):2611–20.

Deng L, Costa F, Blake KJ, Choi S, Chandrabalan A, Yousuf MS, et al. S. aureus drives itch and scratch-induced skin damage through a V8 protease-PAR1 axis. Cell 2023;186(24):5375–93.e25.

Doneddu PE, Borroni R, Ceribelli A, Carta F, Sechi M, Moretti GS, et al. Risk of peripheral neuropathy in patients with psoriasis and psoriatic arthritis. A prospective cohort study. Muscle & Nerve 2024;70(3):371–8.

El-Nour H, Santos A, Nordin M, Jonsson P, Svensson M, Nordlind K, et al. Neuronal changes in psoriasis exacerbation. Journal of the European Academy of Dermatology and Venereology 2009;23(11):1240–5.

Enamorado M, Kulalert W, Han S-J, Rao I, Delaleu J, Link VM, et al. Immunity to the microbiota promotes sensory neuron regeneration. Cell 2023;186(3):607–20.e17.

Fassett MS, Braz JM, Castellanos CA, Salvatierra JJ, Sadeghi M, Yu X, et al. IL-31–dependent neurogenic inflammation restrains cutaneous type 2 immune response in allergic dermatitis. Science Immunology;8(88):eabi6887.

Flayer CH, Kernin IJ, Matatia PR, Zeng X, Yarmolinsky DA, Han C, et al. A γδ T cell–IL-3 axis controls allergic responses through sensory neurons. Nature 2024;634(8033):440–6.

Furue M, Furue K, Tsuji G, Nakahara T. Interleukin-17A and Keratinocytes in Psoriasis. Int J Mol Sci 2020;21(4).

Geherin SA, Lee MH, Wilson RP, Debes GF. Ovine skin-recirculating γδ T cells express IFN-γ and IL-17 and exit tissue independently of CCR7. Veterinary Immunology and Immunopathology 2013;155(1):87–97.

Gray EE, Ramírez-Valle F, Xu Y, Wu S, Wu Z, Karjalainen KE, et al. Deficiency in IL-17-committed Vγ4+ γδ T cells in a spontaneous Sox13-mutant CD45.1+ congenic mouse substrain provides protection from dermatitis. Nature immunology 2013;14(6):584–92.

Grolleau C, Calugareanu A, Demouche S, Nosbaum A, Staumont-Sallé D, Aubert H, et al. IL-4/IL-13 Inhibitors for Atopic Dermatitis Induce Psoriatic Rash Transcriptionally Close to Pustular Psoriasis. J Invest Dermatol 2023;143(5):711–21.e7.

Guo C, Jiang H, Huang C-C, Li F, Olson W, Yang W, et al. Pain and itch coding mechanisms of polymodal sensory neurons. Cell Reports 2023;42(11):113316.

Guo J, Zhang H, Lin W, Lu L, Su J, Chen X. Signaling pathways and targeted therapies for psoriasis. Signal Transduction and Targeted Therapy 2023;8(1):437.

Han L, Ma C, Liu Q, Weng H-J, Cui Y, Tang Z, et al. A subpopulation of nociceptors specifically linked to itch. Nature Neuroscience 2013;16(2):174–82.

Hanč P, Gonzalez RJ, Mazo IB, Wang Y, Lambert T, Ortiz G, et al. Multimodal control of dendritic cell functions by nociceptors. Science;379(6639):eabm5658.

Hao Y, Hao S, Andersen-Nissen E, Mauck WM, III, Zheng S, Butler A, et al. Integrated analysis of multimodal single-cell data. Cell 2021;184(13):3573–87.e29.

Hashimoto T, Sakai K, Sanders KM, Yosipovitch G, Akiyama T. Antipruritic Effects of Janus Kinase Inhibitor Tofacitinib in a Mouse Model of Psoriasis. Acta Derm Venereol 2019;99(3):298–303.

Hau CS, Shimizu T, Tada Y, Kamata M, Takeoka S, Shibata S, et al. The vitamin D 3 analog, maxacalcitol, reduces psoriasiform skin inflammation by inducing regulatory T cells and downregulating IL-23 and IL-17 production. Journal of Dermatological Science 2018;92(2):117–26.

Hu Y, Hu Q, Li Y, Lu L, Xiang Z, Yin Z, et al. gammadelta T cells: origin and fate, subsets, diseases and immunotherapy. Signal Transduct Target Ther 2023;8(1):434.

Ibusuki A, Kawai K, Nitahara-Takeuchi A, Argüello RJ, Kanekura T. TCR signaling and cellular metabolism regulate the capacity of murine epidermal γδ T cells to rapidly produce IL-13 but not IFN-γ. Frontiers in Immunology 2024;Volume 15 - 2024.

Inclan-Rico JM, Napuri CM, Lin C, Hung L-Y, Ferguson AA, Liu X, et al. MrgprA3 neurons drive cutaneous immunity against helminths through selective control of myeloid-derived IL-33. Nature Immunology 2024.

Inclan-Rico JM, Stephenson A, Napuri CM, Rossi HL, Hung L-Y, Pastore CF, et al. TRPV1+ neurons promote cutaneous immunity against Schistosoma mansoni. The Journal of Immunology 2025.

Jean EE, Rossi HL, Hung LY, Inclan-Rico JM, Herbert DR. Myeloid-derived IL-33 drives gammadelta T cell-dependent resistance against cutaneous infection by Strongyloides ratti. J Immunol 2025;214(3):502–15.

Jia J, Li C, Yang J, Wang X, Li R, Luo S, et al. Yes-associated protein promotes the abnormal proliferation of psoriatic keratinocytes via an amphiregulin dependent pathway. Scientific Reports 2018;8(1):14513.

Jin S-P, Koh S-j, Yu D-A, Kim M-W, Yun HT, Lee DH, et al. Imiquimod-applied Interleukin-10 deficient mice better reflects severe and persistent psoriasis with systemic inflammatory state. Experimental Dermatology 2018;27(1):43–9.

Junsuwan N, Likittanasombat S, Chularojanamontri L, Chaiyabutr C, Wongpraparut C, Silpa-archa N. Prevalence and clinical characteristics of pruritus, and the factors significantly associated with high pruritic intensity in patients with psoriasis: a cross-sectional study. Annals of Medicine and Surgery 2023;85(7).

Kashem Sakeen W, Riedl Maureen S, Yao C, Honda Christopher N, Vulchanova L, Kaplan Daniel H. Nociceptive Sensory Fibers Drive Interleukin-23 Production from CD301b^+^ Dermal Dendritic Cells and Drive Protective Cutaneous Immunity. Immunity 2015;43(3):515–26.

Kim HJ, Kim SH, Je JH, Shin DY, Kim DS, Lee M-G. Increased expression of Toll-like receptors 3, 7, 8 and 9 in peripheral blood mononuclear cells in patients with psoriasis. Experimental Dermatology 2016;25(6):485–7.

Kim SJ, Park GH, Kim D, Lee J, Min H, Wall E, et al. Analysis of cellular and behavioral responses to imiquimod reveals a unique itch pathway in transient receptor potential vanilloid 1 (TRPV1)-expressing neurons. Proceedings of the National Academy of Sciences of the United States of America 2011;108(8):3371–6.

Kim T-W, Shim W-H, Kim J-M, Mun J-H, Song M, Kim H-S, et al. Clinical Characteristics of Pruritus in Patients with Scalp Psoriasis and Their Relation with Intraepidermal Nerve Fiber Density. Ann Dermatol 2014;26(6):727–32.

Krishnan S, Prise IE, Wemyss K, Schenck LP, Bridgeman HM, McClure FA, et al. Amphiregulin-producing γδ T cells are vital for safeguarding oral barrier immune homeostasis. Proceedings of the National Academy of Sciences 2018;115(42):10738–43.

Krogstad AL, Lönnroth P, Larson G, Wallin BG. Increased interstitial histamine concentration in the psoriatic plaque. The Journal of investigative dermatology 1997;109(5):632–5.

Lande R, Gregorio J, Facchinetti V, Chatterjee B, Wang Y-H, Homey B, et al. Plasmacytoid dendritic cells sense self-DNA coupled with antimicrobial peptide. Nature 2007;449(7162):564–9.

Liu AW, Zhang YR, Chen C-S, Edwards TN, Ozyaman S, Ramcke T, et al. Scratching promotes allergic inflammation and host defense via neurogenic mast cell activation. Science 2025;387(6733):eadn9390.

Liu Q, Sikand P, Ma C, Tang Z, Han L, Li Z, et al. Mechanisms of Itch Evoked by β-Alanine. The Journal of Neuroscience 2012;32(42):14532–7.

Liu Q, Tang Z, Surdenikova L, Kim S, Patel KN, Kim A, et al. Sensory Neuron-Specific GPCR Mrgprs Are Itch Receptors Mediating Chloroquine-Induced Pruritus. Cell 2009;139(7):1353–65.

Ma F, Plazyo O, Billi AC, Tsoi LC, Xing X, Wasikowski R, et al. Single cell and spatial sequencing define processes by which keratinocytes and fibroblasts amplify inflammatory responses in psoriasis. Nature Communications 2023;14(1):3455.

Madisen L, Zwingman TA, Sunkin SM, Oh SW, Zariwala HA, Gu H, et al. A robust and high-throughput Cre reporting and characterization system for the whole mouse brain. Nature Neuroscience 2010;13(1):133–40.

Mavropoulos A, Varna A, Zafiriou E, Liaskos C, Alexiou I, Roussaki-Schulze A, et al. IL-10 producing Bregs are impaired in psoriatic arthritis and psoriasis and inversely correlate with IL-17- and IFNγ-producing T cells. Clinical Immunology 2017;184:33–41.

McCoy Eric S, Taylor-Blake B, Street Sarah E, Pribisko Alaine L, Zheng J, Zylka Mark J. Peptidergic CGRPα Primary Sensory Neurons Encode Heat and Itch and Tonically Suppress Sensitivity to Cold. Neuron 2013;78(1):138–51.

McKenzie DR, Hart R, Bah N, Ushakov DS, Muñoz-Ruiz M, Feederle R, et al. Normality sensing licenses local T cells for innate-like tissue surveillance. Nature Immunology 2022;23(3):411–22.

Nagel G, Szellas T, Huhn W, Kateriya S, Adeishvili N, Berthold P, et al. Channelrhodopsin-2, a directly light-gated cation-selective membrane channel. Proceedings of the National Academy of Sciences 2003;100(24):13940–5.

Nakao M, Sugaya M, Fujita H, Miyagaki T, Morimura S, Shibata S, et al. TLR2 Deficiency Exacerbates Imiquimod-Induced Psoriasis-Like Skin Inflammation through Decrease in Regulatory T Cells and Impaired IL-10 Production. International journal of molecular sciences 2020. p. 8560.

Narbutt J, Olejniczak I, Sobolewska-Sztychny D, Sysa-Jedrzejowska A, Słowik-Kwiatkowska I, Hawro T, et al. Narrow band ultraviolet B irradiations cause alteration in interleukin-31 serum level in psoriatic patients. Archives of Dermatological Research 2013;305(3):191–5.

Olson W, Abdus-Saboor I, Cui L, Burdge J, Raabe T, Ma M, et al. Sparse genetic tracing reveals regionally specific functional organization of mammalian nociceptors. eLife 2017;6:e29507.

Oprée A, Kress M. Involvement of the Proinflammatory Cytokines Tumor Necrosis Factor-α, IL-1β, and IL-6 But Not IL-8 in the Development of Heat Hyperalgesia: Effects on Heat-Evoked Calcitonin Gene-Related Peptide Release from Rat Skin. The Journal of Neuroscience 2000;20(16):6289–93.

Pantelyushin S, Haak S, Ingold B, Kulig P, Heppner FL, Navarini AA, et al. Rorγt+ innate lymphocytes and γδ T cells initiate psoriasiform plaque formation in mice. The Journal of clinical investigation 2012;122(6):2252–6.

Pergolizzi S, Vaccaro M, Magaudda L, Mondello MR, Arco A, Bramanti P, et al. Immunohistochemical study of epidermal nerve fibres in involved and uninvolved psoriatic skin using confocal laser scanning microscopy. Arch Dermatol Res 1998;290(9):483–9.

Peters C, Kabelitz D, Wesch D. Regulatory functions of gammadelta T cells. Cell Mol Life Sci 2018;75(12):2125–35.

Petersen LJ, Hansen U, Kristensen JK, Nielsen H, Skov PS, Nielsen HJ. Studies on mast cells and histamine release in psoriasis: the effect of ranitidine. Acta Derm Venereol 1998;78(3):190–3.

Que M, Wei Y, Wu S, Liang X, Zhang K, Wu G. Establishment of Psoriasis Mouse Model by Imiquimod. J Vis Exp 2025(220).

Ramirez-Valle F, Gray EE, Cyster JG. Inflammation induces dermal Vgamma4+ gammadeltaT17 memory-like cells that travel to distant skin and accelerate secondary IL-17-driven responses. Proc Natl Acad Sci U S A 2015;112(26):8046–51.

Ramírez-Valle F, Gray EE, Cyster JG. Inflammation induces dermal Vγ4+ γδT17 memory-like cells that travel to distant skin and accelerate secondary IL-17–driven responses. Proceedings of the National Academy of Sciences 2015;112(26):8046–51.

Reich A, Orda A, Wiśnicka B, Szepietowski JC. Plasma concentration of selected neuropeptides in patients suffering from psoriasis. Experimental Dermatology 2007;16(5):421–8.

Reich K, Gräfe A, Vente C, Neumann C, Brück M, Garbe C. Treatment of Psoriasis with Interleukin-10. Journal of Investigative Dermatology 1998;111(6):1235–6.

Rhodes KA, Andrew EM, Newton DJ, Tramonti D, Carding SR. A subset of IL-10-producing γδ T cells protect the liver from Listeria-elicited, CD8+ T cell-mediated injury. European Journal of Immunology 2008;38(8):2274–83.

Riol-Blanco L, Ordovas-Montanes J, Perro M, Naval E, Thiriot A, Alvarez D, et al. Nociceptive sensory neurons drive interleukin-23-mediated psoriasiform skin inflammation. Nature 2014;510(7503):157–61.

Rizzo HL, Kagami S, Phillips KG, Kurtz SE, Jacques SL, Blauvelt A. IL-23–Mediated Psoriasis-Like Epidermal Hyperplasia Is Dependent on IL-17A. The Journal of Immunology 2011;186(3):1495–502.

Rodríguez-Gómez IM, Talker SC, Käser T, Stadler M, Reiter L, Ladinig A, et al. Expression of T-Bet, Eomesodermin, and GATA-3 Correlates With Distinct Phenotypes and Functional Properties in Porcine γδ T Cells. Frontiers in Immunology 2019;Volume 10 - 2019.

Sakai K, Sanders KM, Youssef MR, Yanushefski KM, Jensen L, Yosipovitch G, et al. Mouse model of imiquimod-induced psoriatic itch. PAIN 2016;157(11).

Schwarz A, Philippsen R, Schwarz T. Induction of Regulatory T Cells and Correction of Cytokine Disbalance by Short-Chain Fatty Acids: Implications for Psoriasis Therapy. Journal of Investigative Dermatology 2021;141(1):95–104.e2.

Shallev L, Kopel E, Feiglin A, Leichner GS, Avni D, Sidi Y, et al. Decreased A-to-I RNA editing as a source of keratinocytes’ dsRNA in psoriasis. Rna 2018;24(6):828–40.

Sharif B, Ase AR, Ribeiro-da-Silva A, Séguéla P. Differential Coding of Itch and Pain by a Subpopulation of Primary Afferent Neurons. Neuron 2020;106(6):940–51.e4.

Smith RL, Hébert HL, Massey J, Bowes J, Marzo-Ortega H, Ho P, et al. Association of Toll-like receptor 4 (TLR4) with chronic plaque type psoriasis and psoriatic arthritis. Archives of Dermatological Research 2016;308(3):201–5.

Solinski HJ, Kriegbaum MC, Tseng P-Y, Earnest TW, Gu X, Barik A, et al. Nppb Neurons Are Sensors of Mast Cell-Induced Itch. Cell Reports 2019;26(13):3561–73.e4.

Strid J, Sobolev O, Zafirova B, Polic B, Hayday A. The Intraepithelial T Cell Response to NKG2D-Ligands Links Lymphoid Stress Surveillance to Atopy. Science 2011;334(6060):1293–7.

Sumaria N, Roediger B, Ng LG, Qin J, Pinto R, Cavanagh LL, et al. Cutaneous immunosurveillance by self-renewing dermal γδ T cells. Journal of Experimental Medicine 2011;208(3):505–18.

Taneda K, Tominaga M, Negi O, Tengara S, Kamo A, Ogawa H, et al. Evaluation of epidermal nerve density and opioid receptor levels in psoriatic itch. British Journal of Dermatology 2011;165(2):277–84.

Taniguchi T, Md Mannoor K, Nonaka D, Toma H, Li C, Narita M, et al. A Unique Subset of γδ T Cells Expands and Produces IL-10 in Patients with Naturally Acquired Immunity against Falciparum Malaria. Frontiers in Microbiology 2017;Volume 8 - 2017.

Tseng P-Y, Hoon MA. Oncostatin M can sensitize sensory neurons in inflammatory pruritus. Science translational medicine;13(619):eabe3037.

Ugur M, Kaminski A, Pabst O. Lymph node γδ and αβ CD8+ T cells share migratory properties. Scientific Reports 2018;8(1):8986.

Usoskin D, Furlan A, Islam S, Abdo H, Lönnerberg P, Lou D, et al. Unbiased classification of sensory neuron types by large-scale single-cell RNA sequencing. Nature Neuroscience 2015;18(1):145–53.

Valero-Pacheco N, Tang EK, Massri N, Loia R, Chemerinski A, Wu T, et al. Maternal IL-33 critically regulates tissue remodeling and type 2 immune responses in the uterus during early pregnancy in mice. Proceedings of the National Academy of Sciences 2022;119(35):e2123267119.

van der Fits L, Mourits S, Voerman JSA, Kant M, Boon L, Laman JD, et al. Imiquimod-Induced Psoriasis-Like Skin Inflammation in Mice Is Mediated via the IL-23/IL-17 Axis1. The Journal of Immunology 2009;182(9):5836–45.

Van Gassen S, Callebaut B, Van Helden MJ, Lambrecht BN, Demeester P, Dhaene T, et al. FlowSOM: Using self-organizing maps for visualization and interpretation of cytometry data. Cytometry Part A 2015;87(7):636–45.

Van Rhijn I, Rutten VPMG, Charleston B, Smits M, van Eden W, Koets AP. Massive, sustained γδ T cell migration from the bovine skin in vivo. Journal of Leukocyte Biology 2007;81(4):968–73.

Vrieling M, Santema W, Van Rhijn I, Rutten V, Koets A. γδ T Cell Homing to Skin and Migration to Skin-Draining Lymph Nodes Is CCR7 Independent. The Journal of Immunology 2012;188(2):578–84.

Wang K, Zhao Y, Cao X. Global burden and future trends in psoriasis epidemiology: insights from the global burden of disease study 2019 and predictions to 2030. Archives of Dermatological Research 2024;316(4):114.

Wang L, Su X, Yan J, Wu Q, Xu X, Wang X, et al. Involvement of Mrgprd-expressing nociceptors-recruited spinal mechanisms in nerve injury-induced mechanical allodynia. iScience 2023;26(5):106764.

Xin Q, Zhang W, Yuan S. The Mechanism of the Channel Opening in Channelrhodopsin-2: A Molecular Dynamics Simulation. International Journal of Molecular Sciences2023.

Xing Y, Chen J, Hilley H, Steele H, Yang J, Han L. Molecular Signature of Pruriceptive MrgprA3^+^ Neurons. Journal of Investigative Dermatology 2020;140(10):2041–50.

Yanaba K, Kamata M, Ishiura N, Shibata S, Asano Y, Tada Y, et al. Regulatory B cells suppress imiquimod-induced, psoriasis-like skin inflammation. J Leukoc Biol 2013;94(4):563–73.

Yu H, Nagi SS, Usoskin D, Hu Y, Kupari J, Bouchatta O, et al. Leveraging deep single-soma RNA sequencing to explore the neural basis of human somatosensation. Nature Neuroscience 2024.

Zhang S, Edwards TN, Chaudhri VK, Wu J, Cohen JA, Hirai T, et al. Nonpeptidergic neurons suppress mast cells via glutamate to maintain skin homeostasis. Cell 2021;184(8):2151–66.e16.

Zhao SS, Hyrich K, Yiu Z, Barton A, Bowes J. Genetically Proxied Interleukin-13 Inhibition Is Associated With Risk of Psoriatic Disease: A Mendelian Randomization Study. Arthritis Rheumatol 2024;76(11):1602–10.

Zhou Y, Follansbee T, Wu X, Han D, Yu S, Domocos DT, et al. TRPV1 mediates inflammation and hyperplasia in imiquimod (IMQ)-induced psoriasiform dermatitis (PsD) in mice. J Dermatol Sci 2018;92(3):264–71.

Zhu R, Cai X, Zhou C, Li Y, Zhang X, Li Y, et al. Dermal Vγ(4)(+)T cells enhance the IMQ-induced psoriasis-like skin inflammatidon in re-challenged mice. Am J Transl Res 2017;9(12):5347–60.

Zylka MJ, Rice FL, Anderson DJ. Topographically Distinct Epidermal Nociceptive Circuits Revealed by Axonal Tracers Targeted to <em>Mrgprd</em>. Neuron 2005;45(1):17–25.

