## Supplemental information for "Sensory neurons shape γδ T cell effector programs to control Psoriasiform Inflammation"

### Supplementary Information

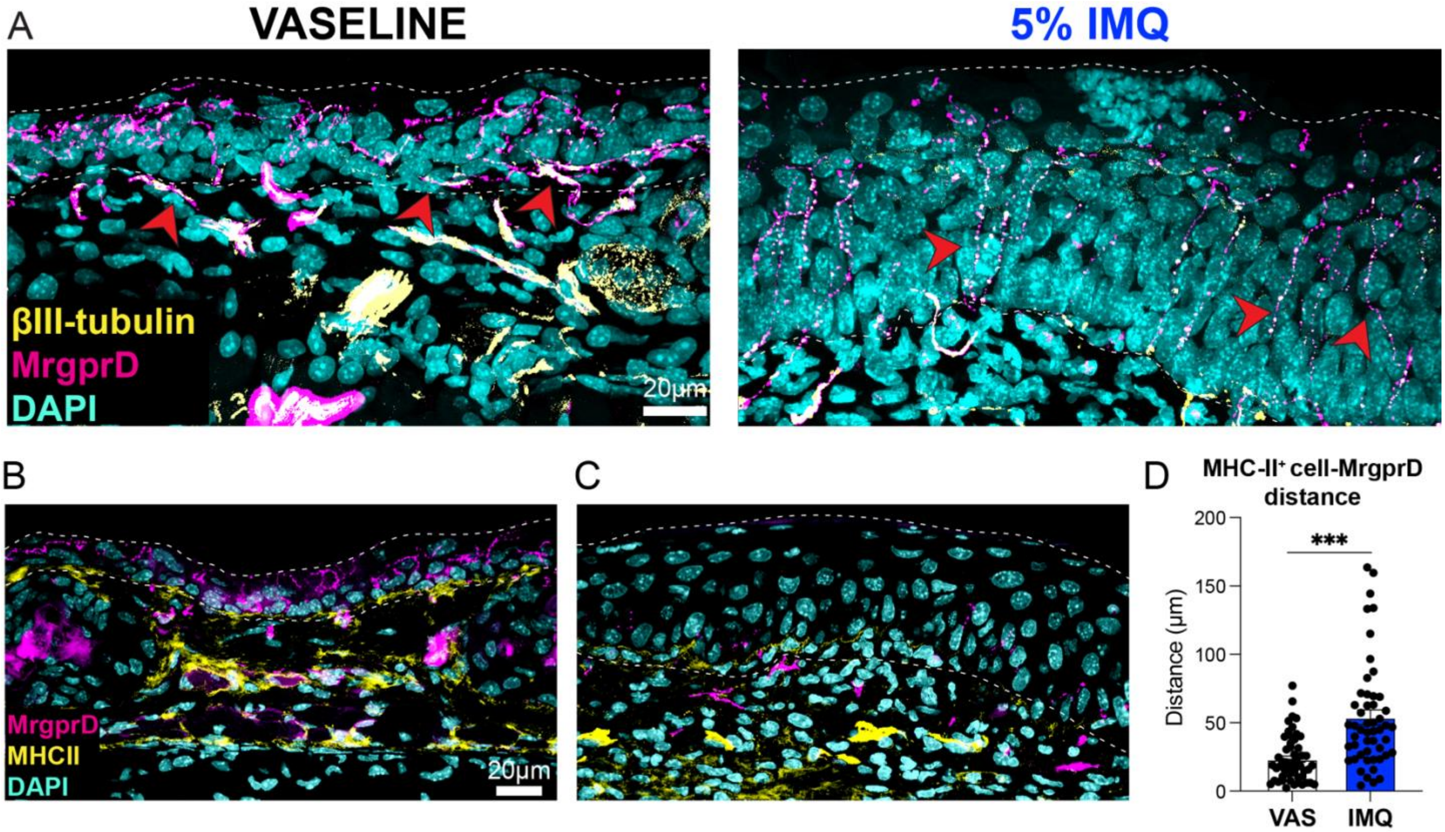

**Supplemental Figure 1. Analysis of MrgprD+ epidermal afferents and antigen-presenting cells (APCs) after IMQ exposure.** (**A**), IFA staining of βIII-tubulin+ MrgprD-tdTomato+ nerve afferents in the epidermis of MrgprD-tdTomato reporter mice treated with Vaseline or 5% IMQ cream. Red arrows indicate co-expression. (**B,C**), IFA staining of epidermal MrgprD-tdTomato+ afferents and MHC-II+ APCs (yellow) in Vaseline- or IMQ-treated mice. (**D**), Mean ± s.e.m (n=51-54) of the distance between APCs and MrgprD+ nerve afferents in the epidermis. *P* values were determined by two-tailed Student's t-test. ***$P<0.001$. Representative of 2 independent experiments, each with ≥3 biological replicates.

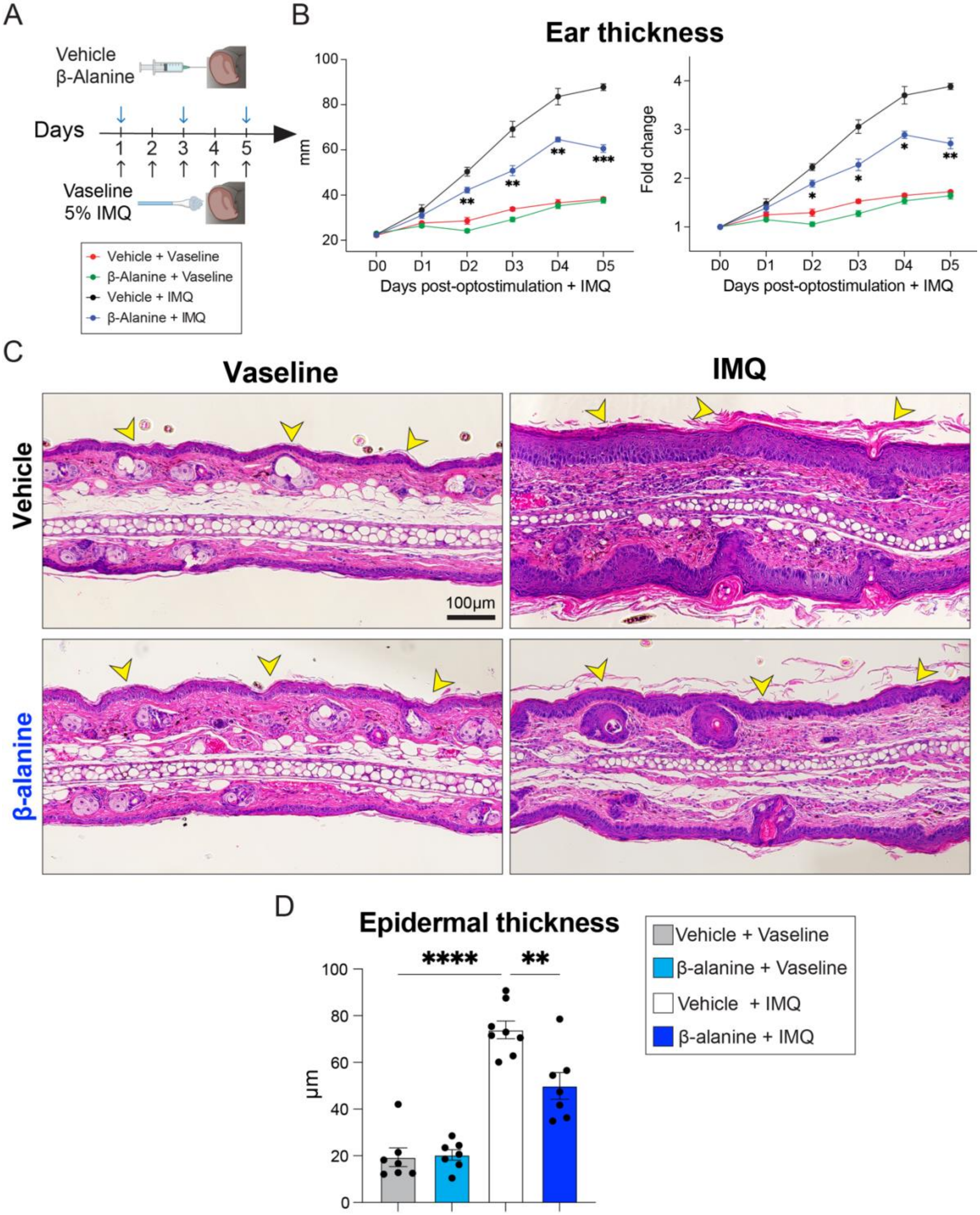

**Supplemental Figure 2. The MrgprD ligand β-alanine ameliorates IMQ-induced psoriatic pathology.** (**A**), Experimental approach for intradermal injection with 20µL of Vehicle (PBS) or 50mM β-alanine in the ear pinnae followed by topical application of Vaseline or 5% IMQ cream. (**B**), Ear thickness was measured daily as shown in A. (**C**), FFPE ear skin sections from mice treated with Vehicle (PBS) or 50mM β-alanine followed by topical application of Vaseline or 5% IMQ cream for 5 days and analyzed by H&E staining. Yellow arrows indicate the epidermis. (**D**), Quantification of epidermal thickness as shown in C. Line or bar graphs depict mean ± s.e.m (n=7-8). *P* values were determined by One-way ANOVA with Bonferroni post hoc correction, or Two-way ANOVA with Tukey's for post hoc test. *P<0.05, **P<0.01, ***P<0.001, ****P<0.0001. Representative of 2-3 independent experiments, each with ≥4 biological replicates.

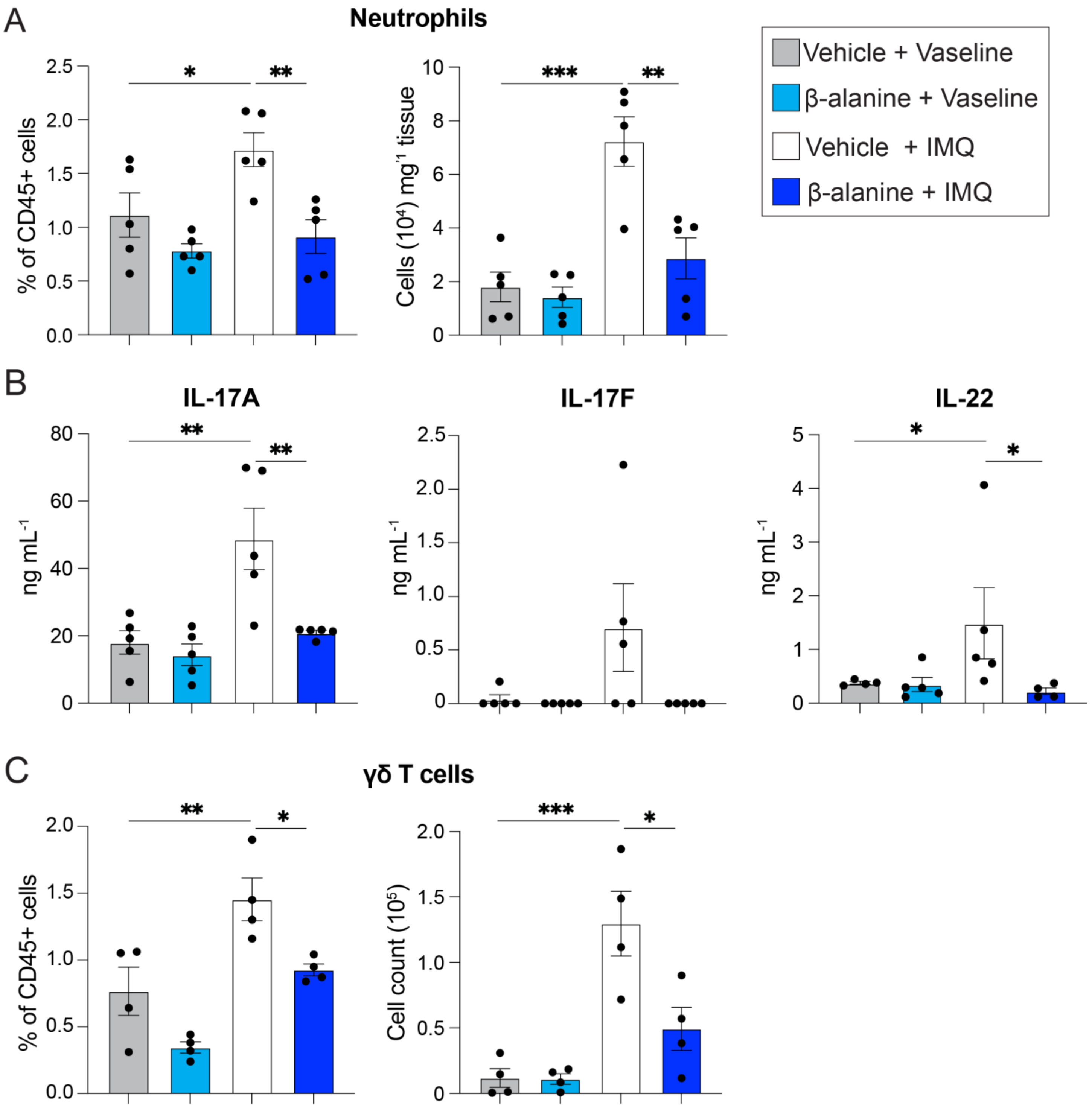

**<u>Supplementary Figure 3. β-alanine treatment suppresses skin IL-17 cytokine responses induced by IMQ.</u>** (**A**), Percentage and absolute numbers of neutrophils (CD45+ CD11b+ Ly6G+ Ly6Cint) of mice treated with Vehicle (PBS) or 50mM β-alanine followed by topical application of Vaseline or 5% IMQ cream. (**B**), Concentration of IL-17A, IL-17F, and IL-22 in cell-free supernatants from sdLNs from Vehicle or β-alanine treated mice during Vaseline or IMQ exposure for 5 days stimulated with αCD3/αCD28 antibodies for 72 h. (**C**), Percentage and absolute numbers of sdLNs γδ T cells (CD45+ CD90+ CD127+ TCRδ+ TCRβ-) from Vehicle or β-alanine-treated mice during Vaseline or IMQ exposure for 5 days. Bar graphs depict mean ± s.e.m (n=4-5). *P* values were determined by One-way ANOVA with Bonferroni post hoc correction. *P<0.05, **P<0.01, ***P<0.001. Representative of 2-3 independent experiments, each with ≥4 biological replicates.

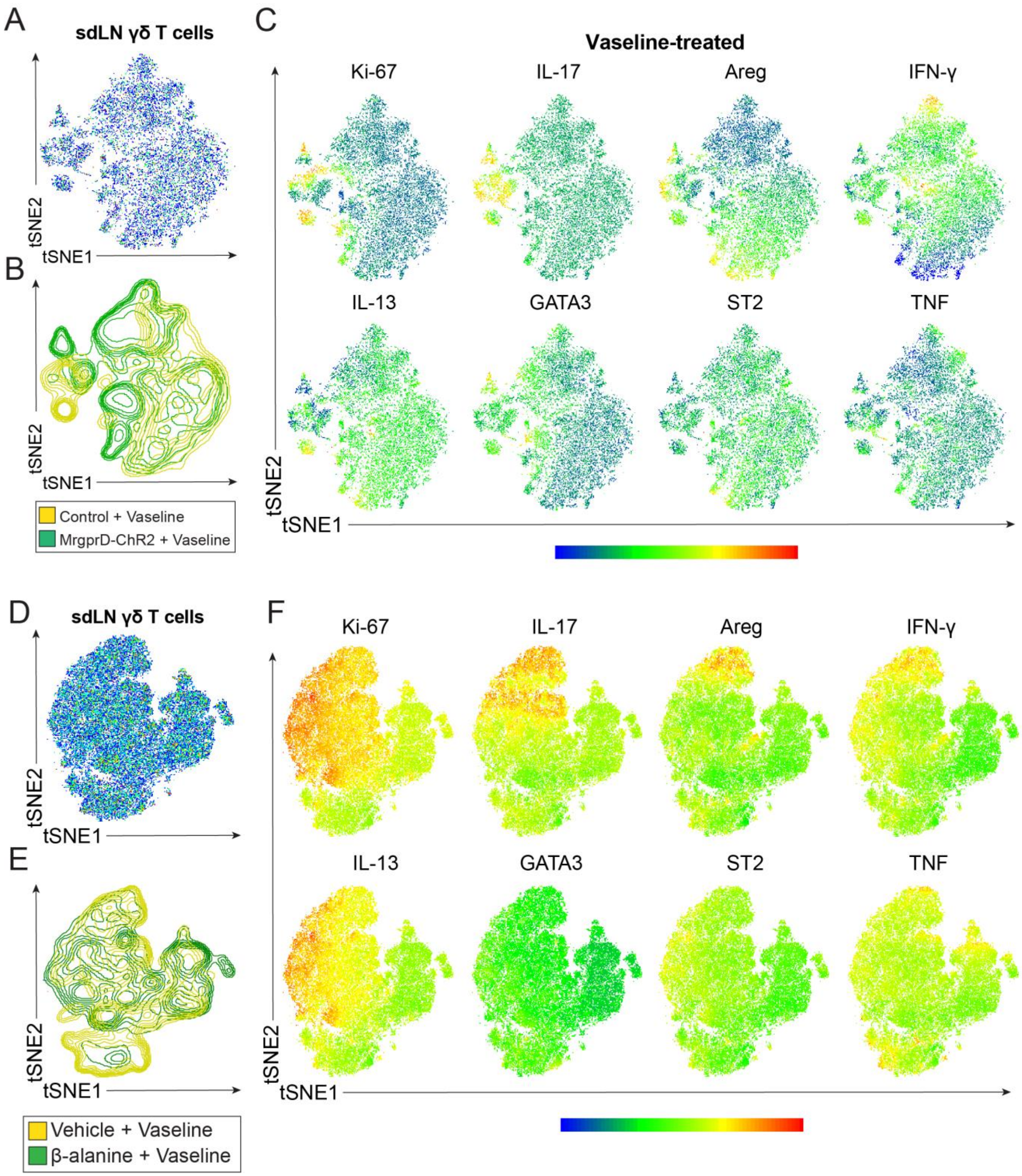

**Supplemental Figure 4. Minor changes of γδ T cells induced by NP1 neuron stimulation in Vaseline-treated mice.** (**A,B**), t-SNE dot plots and counterplots of concatenated γδ T cells from sdLNs of control or MrgprD-ChR2 mice treated for 5 days with Vaseline. (**C**), Expression of cytokines, ST2, GATA-3 and Ki67 in t-SNE dot plots from A. (**D,E**), t-SNE dot plots and counterplots of concatenated γδ T cells from sdLNs of vehicle- and β-alanine-treated mice after 5 days of Vaseline application. (**F**), Expression of cytokines, ST2, GATA-3 and Ki67 in t-SNE dot plots from D.

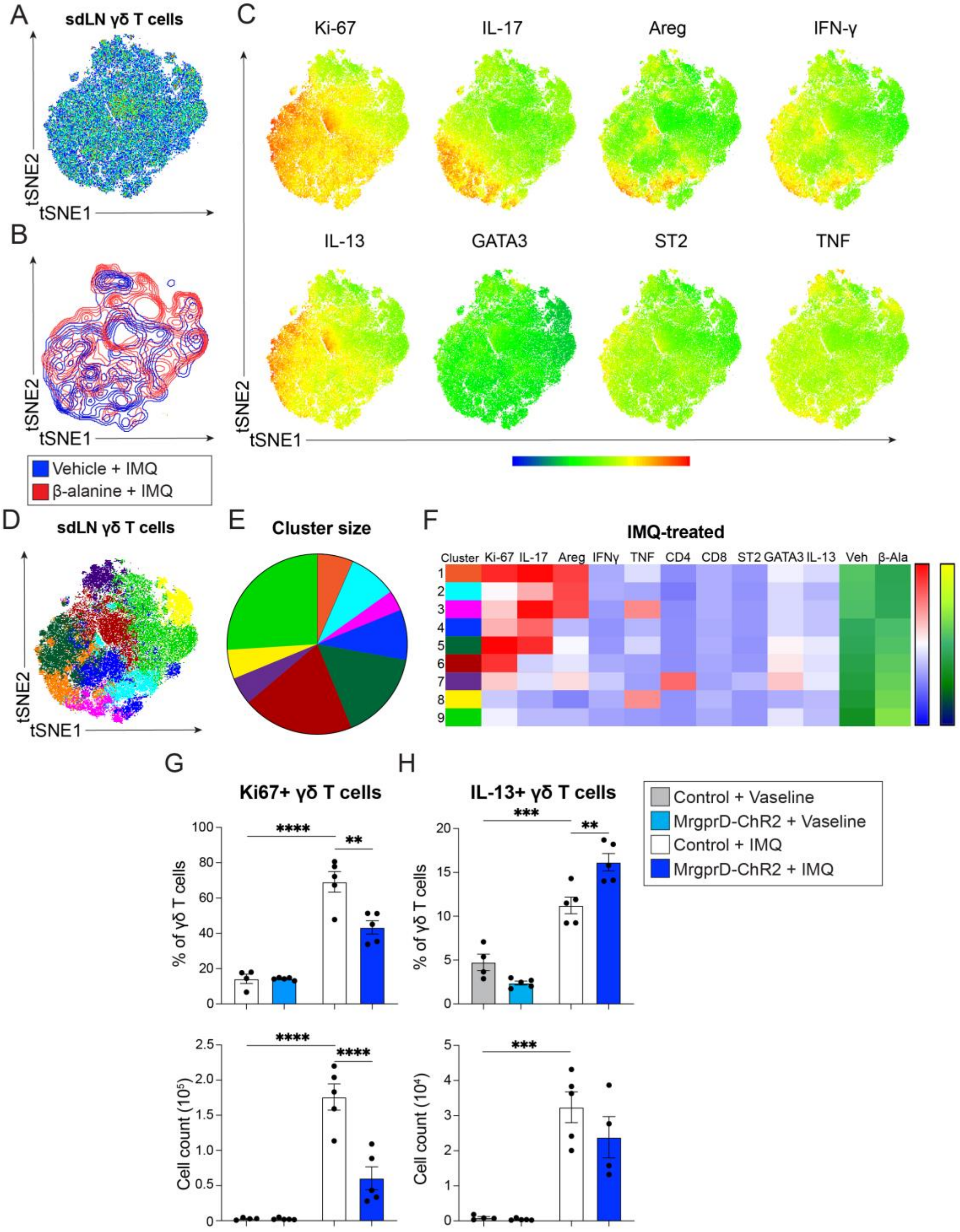

**<u>Supplementary Figure 5. β-alanine treatment regulates the activation profile of γδ T cells during IMQ-induced dermatitis.</u>** (**A,B**), t-SNE dot plots and counterplots of concatenated γδ T cells from sdLNs of mice treated with Vehicle or β-alanine followed by topical application of 5% IMQ cream for 5 days. (**C**), Expression of cytokines, GATA-3 and Ki67 in t-SNE dot plots from A. (**D-F**), FlowSOM analysis, cluster sizes, and heatmaps of γδ T cell clusters from A. (**G,H**), Percentage and absolute numbers of Ki-67$^{+}$ or IL-13$^{+}$ γδ T cells from sdLNs of light-stimulated control or MrgprD-ChR2 mice treated for 5 days with Vaseline or 5% IMQ cream. Bar graphs depict mean ± s.e.m (n=5). *P* values were determined by One-way ANOVA with Bonferroni post hoc correction. *P<0.05, **P<0.01, ***P<0.001, ****P<0.001. Representative of 2-3 independent experiments, each with ≥4 biological replicates.

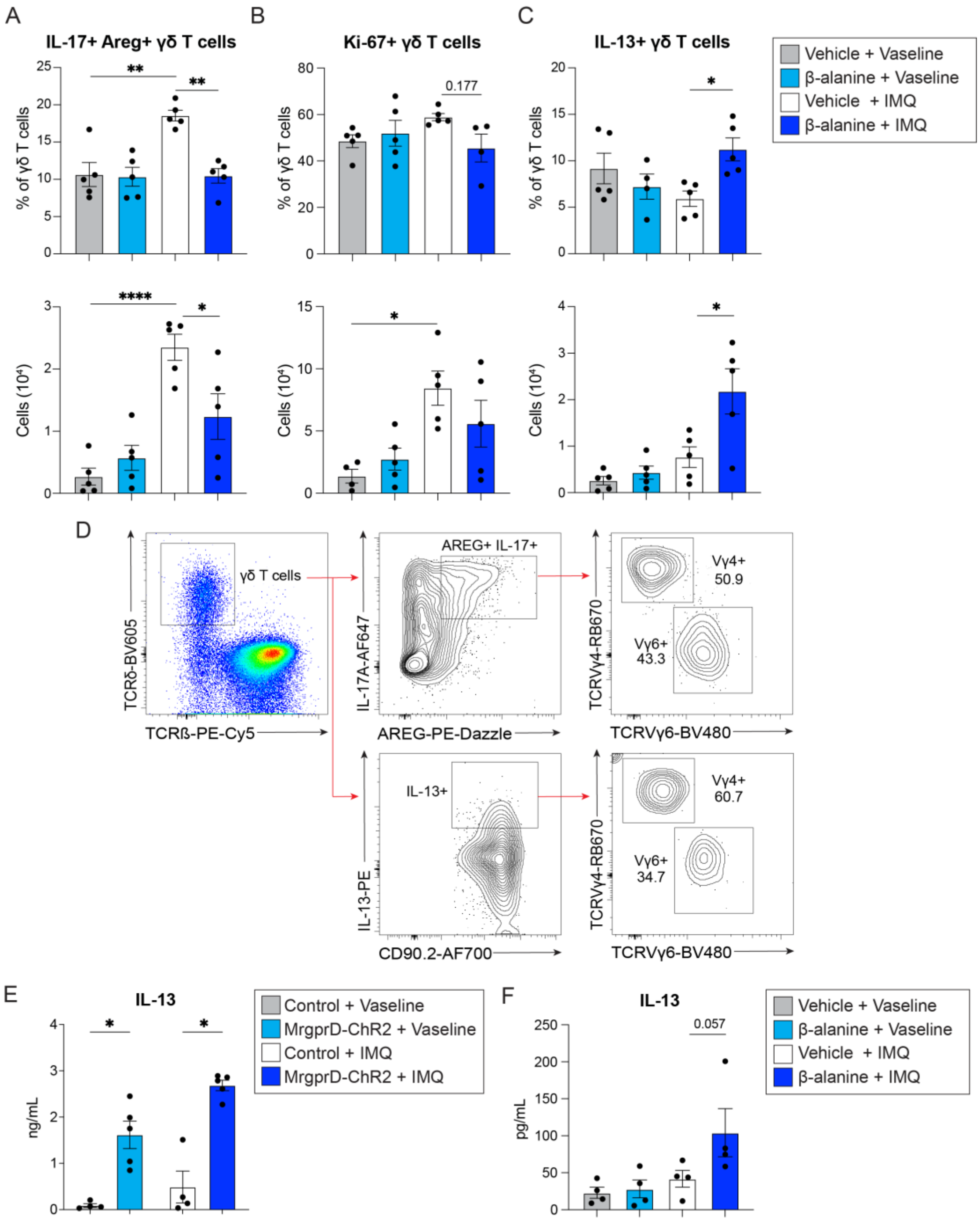

**Supplementary Figure 6. β-alanine treatment reduces IL-17 and stimulates IL-13 expression in γδ T cells during IMQ exposure**. (**A-C**), Percentage and absolute numbers of IL-17$^+$ Areg$^+$, Ki-67$^+$ or IL-13$^+$ γδ T cells from sdLNs of mice treated with Vehicle or β-alanine followed by topical application of Vaseline or IMQ for 5 days. (**D**), Flow cytometric analysis of Vγ4 and Vγ6 populations in IL-17$^+$ Areg$^+$or IL-13$^+$ γδ T cells. (**E**), Concentration of IL-13 in cell-free supernatants from re-stimulated (αCD3/αCD28, 72hrs) sdLNs from control and MrgprD-ChR2 mice or (**F**), vehicle- and β-alanine-treated mice after 5 days of Vaseline or 5% IMQ application. Bar graphs depict mean ± s.e.m (n=4-5). *P* values were determined by One-way ANOVA with Bonferroni post hoc correction. *P<0.05, **P<0.01, ***P<0.001, ****P<0.001. Representative of 2-3 independent experiments, each with ≥4 biological replicates.

| Supplementary Table 1. Resources Table. | | | | | |
|---|---|---|---|---|---|
| REAGENT OR RESOURCE | SOURCE | IDENTIFIER | CLONE | LOTE | Dilution |
| Antibodies | | | | | |
| BUV563™ Rat Anti-Mouse CD45 Antibody | BD Biosciences | Cat# 752412. RRID:AB_2873124 | 30-F11 | 4043831 | 1:250 |
| BUV737™ Rat Anti-Mouse CD11b Antibody | BD Biosciences | Cat# 612801. RRID:AB_2738811 | M1/70 | 3289745 | 1:300 |
| Brilliant Violet 711™ anti-Mouse Ly6G Antibody | Biolegend | Cat# 127643. RRID: AB_2565971 | 1A8 | B406908 | 1:200 |
| Brilliant Violet 785™ anti-Mouse Ly6C Antibody | Biolegend | Cat# 128041. RRID: AB_2565852 | HK1.4 | B406937 | 1:300 |
| BUV395™ Rat Anti-Mouse IL-17 Antibody | BD Biosciences | Cat# 565246. RRID:AB_2722575 | TC11-18H10 | 4117410 | 1:100 |
| BUV737 ™ Rat Anti-Mouse CD4 Antibody | BD Biosciences | Cat# 612844. RRID:AB_2738811 | RM4-5 | 4003189 | 1:200 |
| Brilliant Violet 605™ anti-mouse TCR γ/δ Antibody | Biolegend | Cat# 118129. RRID: AB_2563356 | GL3 | B391618 | 1:200 |
| Alexa Fluor® 700 anti-mouse CD90.2 (Thy1.2) Antibody | Biolegend | Cat# 105319. RRID: AB_493724 | 30-H12 | B364336 | 1:300 |
| Brilliant Violet 785™ anti-mouse IL-33Rα (IL1RL1, ST2) Antibody | Biolegend | Cat# 145321. RRID: AB_2860702 | DIH9 | B409517 | 1:100 |
| Gata-3 Monoclonal Antibody (TWAJ), Alexa Fluor™ 488, eBioscience™ | ThermoFisher | Cat# 53-9966-42. RRID: AB_2574493 | TWAJ | 2619661 | 1:100 |
| Brilliant Violet 711™ anti-mouse CD8a Antibody | Biolegend | Cat# 100747. RRID: AB_11219594 | 53-6.7 | B354913 | 1:200 |
| APC anti-mouse IFN-γ Antibody | Biolegend | Cat# 505810. RRID: AB_315404 | XMG1.2 | B354911 | 1:100 |
| Ki-67 Monoclonal Antibody (SolA15), APC-eFluor™ 780, eBioscience™ | ThermoFisher | Cat# 47-5698-82. RRID: AB_2688065 | SolA15 | 2783661 | 1:200 |
| FOXP3 Monoclonal Antibody (FJK-16s), PE, eBioscience™ | ThermoFisher | Cat#12-5773-82. RRID:AB_465936 | FJK-16s | 2430489 | 1:100 |
| PE/Dazzle™ 594 anti-mouse CD127 (IL-7Rα) Antibody | Biolegend | Cat# 135031. RRID: AB_2564216 | A7R34 | B365389 | 1:100 |
| PE/Cyanine5 anti-mouse TCR β chain Antibody | Biolegend | Cat# 109210. RRID: AB_313433 | H57-597 | B374789 | 1:200 |
| IL-13 Monoclonal Antibody (eBio13A), PE-Cyanine7, eBioscience™ | ThermoFisher | Cat# 25-7133-82. RRID: AB_2573530 | eBio13A | 2764058 | 1:100 |
| BD OptiBuild™ RB670 Mouse Anti-Mouse TCR Vγ4 | BD Biosciences | Cat# 771559. RRID: AB_3693390 | 49.2 | 2023189 | 1:200 |
| BD Horizon™ BV480 Mouse Anti-Mouse TCR Vγ6 | BD Biosciences | Cat# 571001. RRID: AB_3686119 | 1C10-1F7 | 4023484 | 1:200 |
| PerCP/Cyanine5.5 anti-mouse CD3 Antibody | Biolegend | Cat# 100217. RRID: AB_1595597 | 17A2 | N/A | 1:200 |
| PerCP/Cyanine5.5 anti-mouse CD19 Antibody | Biolegend | Cat# 152406. RRID: AB_2629815 | 1D3/CD19 | N/A | 1:200 |
| PerCP/Cyanine5.5 anti-mouse CD11c Antibody | Biolegend | Cat# 117328. RRID: AB_2129641 | N418 | N/A | 1:200 |
| PerCP/Cyanine5.5 anti-mouse CD11b Antibody | Biolegend | Cat# 101228. RRID: AB_893232 | M1/70 | N/A | 1:300 |
| PerCP/Cyanine5.5 anti-mouse NK1.1 Antibody | Biolegend | Cat# 108728. RRID: AB_2132705 | PK136 | N/A | 1:100 |
| PerCP/Cyanine5.5 anti-mouse FcεRIα Antibody | Biolegend | Cat# 134320. RRID: AB_10641135 | MAR-1 | N/A | 1:100 |
| PerCP/Cyanine5.5 anti-mouse CD5 Antibody | Biolegend | Cat# 100624. RRID: AB_2563433 | 53-73.1 | N/A | 1:200 |
| PerCP Cy5.5 anti-mouse Ter119 Antibody | Biolegend | Cat# 116226. RRID: AB_893635 | TER119 | N/A | 1:200 |

| | | | | | |
|---|---|---|---|---|---|
| Keratin 5 Polyclonal Chicken Antibody, Purified | Biolegend | Cat# 905901. RRID: AB_2565054 | poly9059 | B368295 | 1:200 |
| Anti-Cytokeratin 10 antibody | Abcam | Cat# ab76318 | EP1607IHCY | N/A | 1:200 |
| Purified anti-mouse TCR γ/δ Antibody | Biolegend | Cat# 118101. RRID: AB_313826 | GL3 | N/A | 1:100 |
| Anti-MHC Class II antibody | Abcam | Cat# ab139365 | M5/114 | 1063417-12 | 1:200 |
| DAPI Solution | ThermoFisher | Cat# 62248 | N/A | VJ3100051 | 1:1000 |
| Alexa Fluor 594 AffiniPure $F(ab')_2$ Fragment Donkey Anti-Chicken IgY (IgG) (H+L) | Jackson ImmunoResearch Laboratories Inc. | Cat# 703-586-155 | Polyclonal | 165159 | 1:500 |
| AF488 AffiniPure $F(ab')_2$ Fragment Donkey Anti-Rat IgG (H+L) | Jackson ImmunoResearch Laboratories Inc. | Cat# 712-166-150 | Polyclonal | 169156 | 1:500 |
| AF647 AffiniPure $F(ab')_2$ Fragment Donkey Anti-Rabbit IgG (H+L) | Jackson ImmunoResearch Laboratories Inc. | Cat# 711-166-152 | Polyclonal | 167391 | 1:500 |
| Biotin anti-Tubulin β3 (TUBB3) Antibody | BioLegend | Cat#: 801212 | TUBB3 | N/A | 1:100 |
| Cy3 Streptavidin | Jackson ImmunoResearch Laboratories Inc. | Cat# 016-160-084 | Polyclonal | 159372 | 1:200 |
| Alexa Fluor 488 AffiniPure $F(ab')_2$ Fragment Donkey Anti-Chicken IgY (IgG) (H+L) | Jackson ImmunoResearch Laboratories Inc. | Cat# 703-545-155 | Polyclonal | 154923 | 1:500 |
| InVivoMAb anti-mouse IL-10R (CD210) | Bioxcell | Catalog #BE0050 | 1B1.3A | N/A | N/A |
| InVivoMAb rat IgG1 isotype control, anti-horseradish peroxidase | Bioxcell | Catalog #BE0088 | HRPN | N/A | N/A |
| Chemicals, peptides, and recombinant proteins | | | | | |
| Collagenase from *Clostridium histolyticum* | Sigma-Aldrich | Cat# C7657 | | | |
| Hyaluronidase from bovine testes | Sigma-Aldrich | Cat# H3506 | | | |
| DNase I from bovine pancreas | Sigma-Aldrich | Cat# 11284932001 | | | |
| Cell Activation Cocktail (with Brefeldin A) | Biolegend | Cat# 423304 | | | |
| Collagenase A from *Clostridium histolyticum* | Sigma-Aldrich | Cat# 10103578001 | | | |
| Collagenase D | Roche | Cat# 11088866001 | | | |
| Dispase II | Sigma-Aldrich | Cat# D4693 | | | |
| β-Alanine | Sigma-Aldrich | Cat# 146064 | | | |
| eBioscience™ Foxp3 / Transcription Factor Staining Buffer Set | ThermoFisher | Cat# 00-5523-00 | | | |
| DMEM/F-12, powder | ThermoFisher | Cat# 12500062 | | | |
| RPMI 1640 Medium, powder | ThermoFisher | Cat# 31800-022 | | | |
| 16% Paraformaldehyde (formaldehyde) aqueous solution | Electron Microscopy Sciences | Cat#15710-S | | | |
| Normal donkey serum | ThermoFisher | Cat# NC9624464 | | | |
| Fluoroshield Histology Mounting Medium | Sigma-Aldrich | Cat# F6182 | | | |
| Commercial Assays | | | | | |
| IL-17A (homodimer) Mouse Uncoated ELISA Kit | ThermoFisher | Cat# 88-7371-88 | | | |
| Mouse IL-17AF (heterodimer) Uncoated ELISA Kit | ThermoFisher | Cat# 88-8711-88 | | | |
| Mouse IL-22 Uncoated ELISA Kit | ThermoFisher | Cat# 88-7422-88 | | | |
| Mouse IL-13 Uncoated ELISA Kit | ThermoFisher | Cat# 88-7137-88 | | | |
| Mouse IL-10 Uncoated ELISA Kit | ThermoFisher | Cat# 88-7105-88 | | | |
| LIVE/DEAD™ Fixable Aqua Dead Cell Stain Kit, for 405 nm excitation | ThermoFisher | Cat# L34966 | | | |

| | | |
|---|---|---|
| Instruments and equipment | | |
| Symphony A3 Lite | BD Biosciences | N/A |
| MITUTOYO Dial Thickness Gauge | Grainger | Part # 6NRC7 |
| 473nm DPSS Laser Nd:YAG | SLOC lasers | Model BL473T8-100FC |
| Power source | SLOC lasers | Model ADR-800A |
| 1x2 MM Fiber Optic Coupler | Thor Labs | Part# TM200FS1B |
| Power and energy meter | Thor Labs | Part# PM100D |
| 10 MHz Dual Channel Function/Arbitrary Waveform Generator | BX Precision | Part# 4053B |
| Leica DM6000 | Leica | N/A |
| Leica TCS SP8 WLL Confocal | Leica | N/A |
| Software and algorithms | | |
| Flowjo 10.8.1 | Flowjo | N/A |
| Prism 10 | Graph Pad | N/A |
| Leica Application Suite X Version 5.1.0.25446 | Leica | N/A |
